# Self-regulation of single-stranded DNA wrapping dynamics by *E. coli* SSB promotes both stable binding and rapid dissociation

**DOI:** 10.1101/2019.12.20.885368

**Authors:** M. Nabuan Naufer, Michael Morse, Guðfríður Björg Möller, James McIsaac, Ioulia Rouzina, Penny J. Beuning, Mark C. Williams

## Abstract

*E. coli* SSB (*Ec*SSB) is a model protein for studying functions of single-stranded DNA (ssDNA) binding proteins (SSBs), which are critical in genome maintenance. *Ec*SSB forms homotetramers that wrap ssDNA in multiple conformations in order to protect these transiently formed regions during processes such as replication and repair. Using optical tweezers, we measure the binding and wrapping of a single long ssDNA substrate under various conditions and free protein concentrations. We show that *Ec*SSB binds in a biphasic manner, where initial wrapping events are followed by unwrapping events as protein density on the substrate passes a critical saturation. Increasing free *Ec*SSB concentrations increase the fraction of *Ec*SSBs in less-wrapped conformations, including a previously uncharacterized *Ec*SSB_8_ bound state in which ∼8 nucleotides of ssDNA are bound by a single domain of the tetramer with minimal substrate deformation. When the ssDNA is over-saturated with *Ec*SSB, stimulated dissociation rapidly removes excess *Ec*SSB, leaving an array of stably-wrapped *Ec*SSB-ssDNA complexes. We develop a multi-step kinetic model in which *Ec*SSB tetramers transition through multiple wrapped conformations which are regulated through nearest neighbor interactions and ssDNA occupancy. These results provide a mechanism through which otherwise stably bound and wrapped *Ec*SSB tetramers can be rapidly removed from an ssDNA substrate to allow for DNA maintenance and replication functions while still fully protecting ssDNA over a wide range of protein concentrations.

## Introduction

Single-stranded DNA binding proteins (SSBs) rapidly sequester and protect transiently formed single-stranded DNA (ssDNA) segments during genome maintenance (1-12). They exhibit high affinity ssDNA binding and may also play regulatory roles by interacting with other proteins involved in genome maintenance (13, 14). The SSB from *E. coli* (*Ec*SSB) is a model SSB that has been extensively studied.

Monomeric *Ec*SSB is 19 kDa, consists of an oligonucleotide binding (OB) domain-containing N-terminal domain, and a C-terminal domain with a conserved 9-amino acidic tip, and a poorly conserved intrinsically disordered linker (IDL) (15-19). The N-terminal OB domain mediates both inter-protein interactions to form tetramers (which is referred to as *Ec*SSB henceforth), as well as high-affinity DNA binding. *Ec*SSB was shown to exhibit high cooperativity in certain ssDNA binding conformations, which is eliminated by truncating or replacing the IDL or TIP, as well as by mutating the “bridge interface” that links adjacent SSB tetramers through an evolutionarily conserved surface near the ssDNA-binding site (2, 6, 8, 12, 18, 20-22). *Ec*SSB can bind ssDNA with multiple conformations that wrap the ssDNA substrates to different degrees (8, 23-26). The distinct binding modes of *Ec*SSB are identified based on the number of nucleotides (n) occluded by the tetramer upon binding to ssDNA. Solution conditions such as the salt composition and concentration, protein density, as well as template tension have been shown to affect the stability of these distinct binding modes (8, 23-27). Importantly, high cooperativity of binding appears to be typical of the low-salt *Ec*SSB-ssDNA complexes (< 20 mM NaCl, < 1 mM MgCl_2_), when only two out of the four OB sites of the EcSSB tetramer are associated with ssDNA (28). Moreover, it appears that the EcSSB mutants that lack cooperative behavior are fully functional for replication in cells and are able to complement deletion of the *ssb* gene in *E. coli* (22). Thus far, four stable or semi-stable modes, *Ec*SSB_17_, *Ec*SSB_35_, *Ec*SSB_56_, *Ec*SSB_65_ have been identified and directly observed on a stretched ssDNA substrate (27). X-ray crystallographic structural studies revealed a model for the *Ec*SSB_65_ binding topology in which the ssDNA is fully wrapped through the association of all four *Ec*SSB subunits (16). However, the precise topologies of the other binding modes are not completely understood to date.

Recent single molecule analyses have greatly enhanced the understanding of the binding dynamics of these distinct modes (18, 20). These studies have revealed the dynamic equilibrium between well-defined *Ec*SSB functional and structural states (29), and the ability of the tetramer to diffuse quickly along the ssDNA substrate while maintaining its wrapped conformation (30). Nevertheless, several longstanding questions on *Ec*SSB function remain ambiguous, especially with respect to its collective binding dynamics and kinetics. To this end, using optical tweezers, we directly probe the collective binding dynamics of *Ec*SSB to a long ssDNA substrate, especially under conditions that result in protein-saturation of the substrate. For the first time, we investigate the collective binding kinetics and interconversion dynamics of *Ec*SSB with high spatial and temporal resolution. This allows us to characterize a minimally bound *Ec*SSB state that does not wrap ssDNA, which is consistent with only one domain of the tetramer bound to the substrate. We also identify a critical point of protein saturation, above which *Ec*SSB tetramers bind in a competitive fashion, destabilizing the wrapping and binding of their neighbors. These interactions are critical to the seemingly paradoxical function of *Ec*SSB. High affinity and stable binding and its ability to occupy up to 65 nt of ssDNA per tetramer allow *Ec*SSB to fully protect long stretches of ssDNA even under conditions of low free protein concentration. However, during DNA processing events, *Ec*SSB must be rapidly removed as the ssDNA segment shrinks in length. Based on the results from this study, we propose a mechanism for rapid self-regulation of *Ec*SSB density to continuously provide optimal ssDNA coverage during genomic maintenance.

## Results

### Competitive ssDNA binding assay for *Ec*SSB

To characterize the collective ssDNA binding and wrapping kinetics of *Ec*SSB, we generated an 8.1 knt long ssDNA substrate in an optical tweezers system (Fig. 1A). The ssDNA was then stretched, held, and maintained at a tension of 12 pN via a force feedback module while in a protein-free buffer (50 mM Na^+^, 10 mM Hepes, pH 7.5, unless otherwise stated). The buffer surrounding the ssDNA molecule was then rapidly exchanged (<1 s) with a solution containing a fixed *Ec*SSB concentration (Fig. 1B). While the tension along the ssDNA is maintained, the binding of *Ec*SSB to the ssDNA results in a change in ssDNA extension. We observed a biphasic binding profile at saturating *Ec*SSB concentrations (≥1 nM) wherein a rapid shortening of the ssDNA was followed by a slower elongation that equilibrates to an extension less than that of a protein-free ssDNA molecule. Because *Ec*SSB is known to wrap the ssDNA in different modes with varying degrees of ssDNA compaction, the biphasic profile indicates that while *Ec*SSB rapidly wraps the ssDNA after initial binding, the degree of wrapping decreases as more *Ec*SSB saturates the ssDNA substrate (oversaturation). Both the initial rapid ssDNA shortening, and its subsequent partial recovery of extension occur over a longer timescale as the protein concentration is decreased (Fig. 1C). At low enough concentration (∼0.1 nM), the second phase disappears completely, and the extension decreases at a single exponential rate. Additionally, the amplitude of the final, equilibrium change in ssDNA extension decreases as the free *Ec*SSB concentration in solution is increased, which is again consistent with the fact that *Ec*SSB wraps the ssDNA to a lesser degree when the substrate becomes oversaturated.

**Figure 1:**
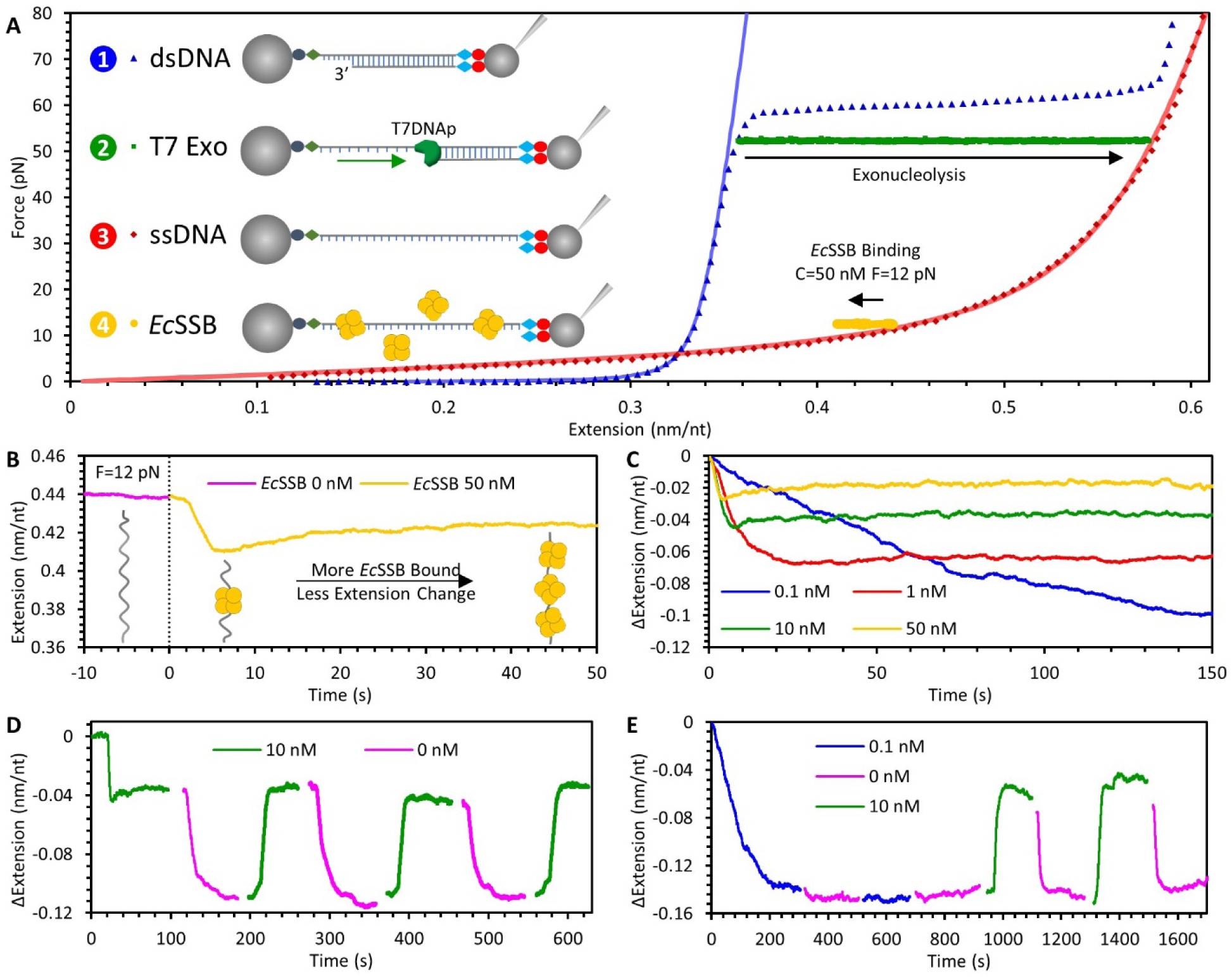
Experimental procedure to measure *Ec*SSB-ssDNA binding dynamics. (A) An 8.1 kbp dsDNA with a recessed 3′ end is tethered between two functionalized beads (step 1, blue). One bead is held by a glass micropipette tip which is moved by a piezo electric stage to extend the DNA. The other bead is held in a stationary dual beam optical trap, the deflection of which measures the force acting on the ssDNA substrate. The dsDNA is incubated with T7 DNA polymerase and held at 50 pN to trigger exonucleolysis to digest the bottom strand (step 2, green), resulting in a long ssDNA molecule (step 3, red). The ssDNA is then held at a constant force and then incubated with varying concentrations of *Ec*SSB (step 4, yellow). Force-extension curves for dsDNA and ssDNA are fit to the WLC and FJC polymer models, respectively. T7 polymerase strand digestion is registered as an increase in the DNA extended length while held at constant force. *Ec*SSB binding results in a decrease in DNA extension. (B) The extension of ssDNA during *Ec*SSB incubation is plotted as a function of time. The ssDNA extension at equilibrium is shorter than bare ssDNA, but longer than the minimum extension achieved immediately after the introduction of *Ec*SSB. (C) Reduction in the *Ec*SSB concentration in the solution increases the net extension change of the *Ec*SSB-ssDNA complex. (D) *Ec*SSB concentration jump experiments showing that removal of the free *Ec*SSB in the solution after initial incubation, results in an extension decrease that stably equilibrates (∼100 s) on a maximally wrapped conformation. Re-introducing *Ec*SSB solution to this equillibriated complex results in an increase in the complex extension, and oscillations between these two extension values are repeatable through changes in free protein concentration. (E) When ssDNA is incubated with low *Ec*SSB concentration, there is no additional change in extension associated with the removal of free protein. Increasing *Ec*SSB concentration, however, still increases the ssDNA’s extension change.

We next measured how *Ec*SSB already bound to the ssDNA substrate reacts to changes in free protein concentration. For each initial *Ec*SSB concentration, after the ssDNA-*Ec*SSB complex reached an equilibrated length, free protein was rapidly removed from the flow cell (Fig. 1D). Interestingly, this resulted in a sudden decrease in ssDNA extension, which was then stable over the timescale of our observation (100s of seconds). In the absence of free protein, the compaction of the ssDNA must result from the *Ec*SSB interconversion to higher order wrapped states that stably equilibrate, as complete dissociation would present as an extension increase that approaches the initial extension of the bare ssDNA. We then reintroduced free protein into the sample, which increased the ssDNA extension back to same equilibrium value achieved during the first incubation. We further show this entire process of adding and removing *Ec*SSB from the sample is repeatable over many cycles, with the ssDNA extension adjusting accordingly. However, when we first incubated the ssDNA with 0.1 nM *Ec*SSB, where the ssDNA extension decreases during incubation is maximized and no biphasic extension increase is observed, removing free protein did not change the ssDNA-*Ec*SSB complex extension (Fig. 1E). Finally, later increasing the free protein concentration did increase the extension, indicating the increases and decreases in ssDNA extension when free *Ec*SSB concentration is changed are fully reversible and the ssDNA-*Ec*SSB complex will reliably equilibrate to a set length based on the current free protein conditions, without regard to previous conditions.

Trying to infer the wrapping kinetics of many *Ec*SSBs on a single ssDNA substrate is greatly complicated by the multiple modes of EcSSB wrapping. However, Suksombat et al. (27), showed that *Ec*SSB_17_ is the stable wrapped state at ssDNA tensions >8 pN. We specifically perform these experiments at 12 pN to inhibit higher order wrapping states, reducing the complexity of this system to a potentially solvable process. Suksombat et al. measured that a single EcSSB in the 17 nt wrapping state reduces the extension of an ssDNA substrate held at 12 pN by ∼5 nm. We confirm this value, by repeating our incubation experiments at a very low *Ec*SSB concentration of 50 pM (Fig. S1), which allows us to temporally resolve individual *Ec*SSB tetramers wrapping and unwrapping on ssDNA as sudden decreases and increases of ssDNA extension with an average amplitude of 5 nm. Moreover, at a lower ssDNA tension of 7 pN, where both the 17 and 35 nt EcSSB wrapped states were observed previously (27), we observe distribution of wrapping event amplitudes consistent with multiple wrapping transitions being present (Fig. S2). Thus, at 12 pN tension, we can characterize our competitive binding assays based on how many *Ec*SSB tetramers are in the 17 nt wrapped state over time and as a function of protein concentration. The inhibition of *Ec*SSB wrapping that we observe at high protein concentration, however, suggests that *Ec*SSB can bind ssDNA in some unwrapped conformation that sterically prohibits other bound proteins from wrapping. As we will show, this bound but unwrapped state, which has been previously uncharacterized, is consistent with a single domain of the *Ec*SSB tetramer binding ∼8 nt of ssDNA substrate (which we will denote as *Ec*SSB_8_), as opposed to wrapped *Ec*SSB_17_, in which two domains bind in order to wrap 17 nt of ssDNA substrate (27).

### General two-step kinetic model for competitive binding dynamics

Based on the observed concentration-dependent collective binding dynamics at 12 pN, we present a generalized two-step kinetic model to describe the competitive *Ec*SSB binding to ssDNA (Fig. 2A):

**Figure 2:**
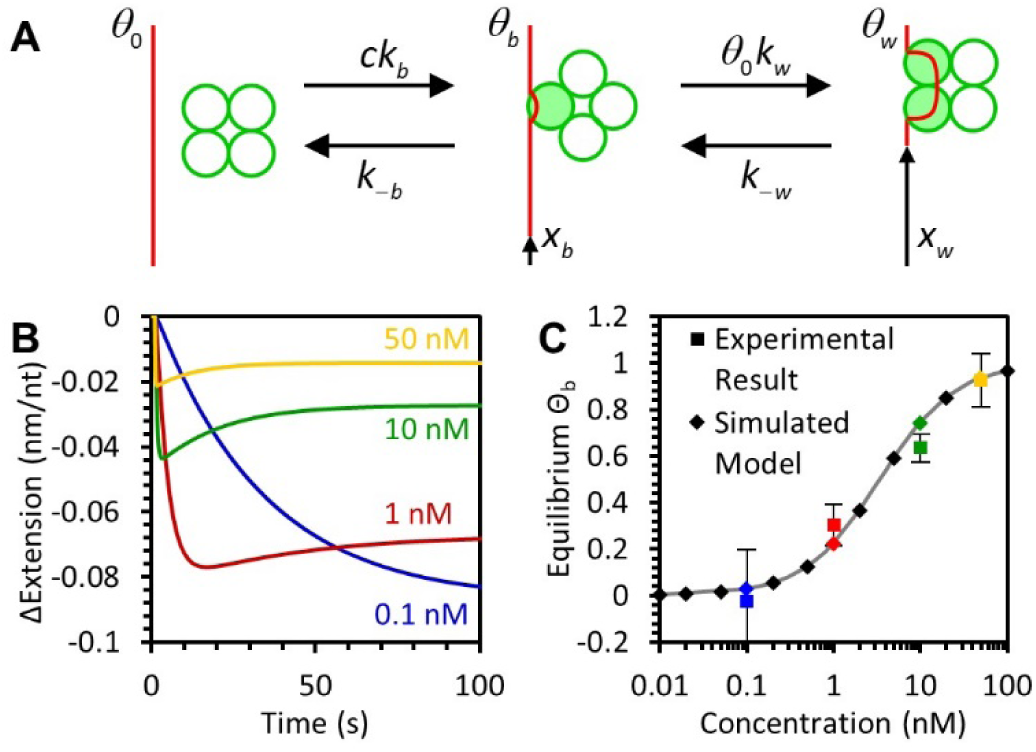
General model of two step binding and wrapping of ssDNA by *Ec*SSB. (A) The protein binds and wraps ssDNA in two distinct steps. First, *Ec*SSB bimolecularly binds to ssDNA in which the on rate is proportional to the protein concentration and dissociation rate is a constant. In the second step, bound protein interconverts between wrapped and unwrapped conformations. The wrapping rate is proportional to the fraction of protein-free ssDNA whereas the unwrapping rate is proportional to the fraction of ssDNA that is occupied by bound but unwrapped protein. The ssDNA extension reduction due an unwrapped *Ec*SSB of the ssDNA is small but measurable, while the wrapped state significantly reduces the ssDNA extension. (B) Simulation of the two-step binding model reproduces the biphasic extension-time profiles that are consistent with experimental results. Substituting the fundamental kinetic rates as estimated from the observed rates in the proposed model reproduces the experimentally observed concentration dependences (Fig. 1C). (C) The fraction of ssDNA-bound *Ec*SSB in the unwrapped state (*Ɵ*_b_) upon reaching equilibrium is predicted by the model (diamonds) and follows the shape of a standard binding isotherm (gray line) where the two states are equally occupied at 3 nM *Ec*SSB. These results agree with experimental data (squares), in which the equilibrium ssDNA extension change is converted into values of *Ɵ*_b_ and *Ɵ*_w_ using the corresponding reductions in ssDNA extension due to each state, X_b_ and X_w_. The solid line represents fit to the free-energy dependence of the binding fractions as derived in the supplemental information.

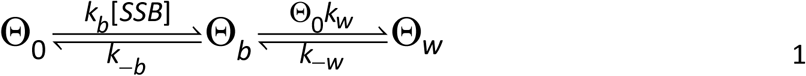

Here, Ɵ_0_, Ɵ_b_, and Ɵ_w_ are the fractions of ssDNA substrate that are protein free, occupied by bound but not wrapped *Ec*SSB_8_, and occupied by wrapped *Ec*SSB_17_, respectively. At 12 pN, we refer specifically to *Ec*SSB_17_ as the wrapped state, though more steps could be added to the right side of this reaction to represent higher order wrapped states accessible at lower forces. *k*_b_ and *k*_-b_ are the bimolecular association and dissociation rates, respectively. *k*_w_ and *k*_-w_ are effective wrapping and unwrapping rates of a single *Ec*SSB, respectively. Additionally, a single bound or wrapped *Ec*SSB reduces ssDNA extension by a value of Δ*x*_b_ or Δ*x*_*w*_, respectively (see Supplemental Information). Given the correct set of parameters (derived in detail in the following sections), this model accurately reproduces the concentration-dependent biphasic binding profiles we observe experimentally (Fig. 2B). Furthermore, the model accurately predicts the equilibrium balance of *Ec*SSB in either the bound or wrapped states over wide range of EcSSB concentrations (Fig 2C). While *Ec*SSB preferentially wraps the ssDNA at low protein concentrations, the wrapped states becomes less stable as the concentration is increased.

### Concentration dependent interconversion of *Ec*SSB states

To quantify the interconversion dynamics of *Ec*SSB, we measure the amplitude of extension change (ΔX) associated with each phase of our binding experiments (Fig. 3A). Each phase is defined by the primary processes responsible for the observed change in the ssDNA extension. First, when *Ec*SSB is initially introduced, the ssDNA shortens as *Ec*SSB binds then wraps the ssDNA (Fig. 3A blue line, ΔX_b,w_), which we denote as the bind-wrap transition. Second, as the ssDNA becomes oversaturated, *Ec*SSB cannot wrap any further, and instead the ssDNA elongates as *Ec*SSB starts to unwrap to accommodate more binding, which we define as the bind-unwrap transition (red line, ΔX_b,-w_). Third, with free protein removed from the solution, no further binding can occur, but some *Ec*SSB dissociates allowing further wrapping of bound *Ec*SSB resulting in ssDNA shortening (green line, ΔX_-b,w_), and this process is referred to as the unbind-wrap transition. Finally, reintroducing free protein once again elongates ssDNA by forcing *Ec*SSB to unwrap to accommodate more protein (yellow line, ΔX_b,-w_). We average the results from three or more independent experiments for each *Ec*SSB concentration (Fig. 3B), showing several significant trends. The amount of ssDNA contraction (and underlying *Ec*SSB wrapping) decreases with increasing *Ec*SSB concentration. However, regardless of the initial *Ec*SSB concentration, the subsequent ssDNA extension upon removal of free protein (ΔX_-b,w_) converges at ∼0.08 nm/nt, which is the same as the net extension change observed with single-phase binding at [*Ec*SSB]≤0.1 nM (Fig. 1C, blue line). Thus, when free *Ec*SSB is scarce, the *Ec*SSB-ssDNA complex reproducibly returns to the same stable equilibrium state, in which excess *Ec*SSB dissociates, and the rest remains stably wrapped. Thus, we designate this net extension as the characteristic extension change associated with the wrapped state (Ɵ_w_). This result also compares favorably with measurements of a single isolated *Ec*SSB tetramer, which decreases the extension of a 65 nt long ssDNA segment by 5 nm, for a normalized compaction of 0.078 nm/nt. Therefore, the *Ec*SSB-ssDNA complex is most stable when the total number of bound tetramers does not exceed one per every ∼70 nt of ssDNA. In contrast, at the highest measured *Ec*SSB concentration, the wrapped state is destabilized, and the *Ec*SSB-ssDNA complex exhibits a small, but non-zero extension change. We therefore associate this ∼0.02 nm/nt extension change with the bound but unwrapped state of *Ec*SSB.

**Figure 3:**
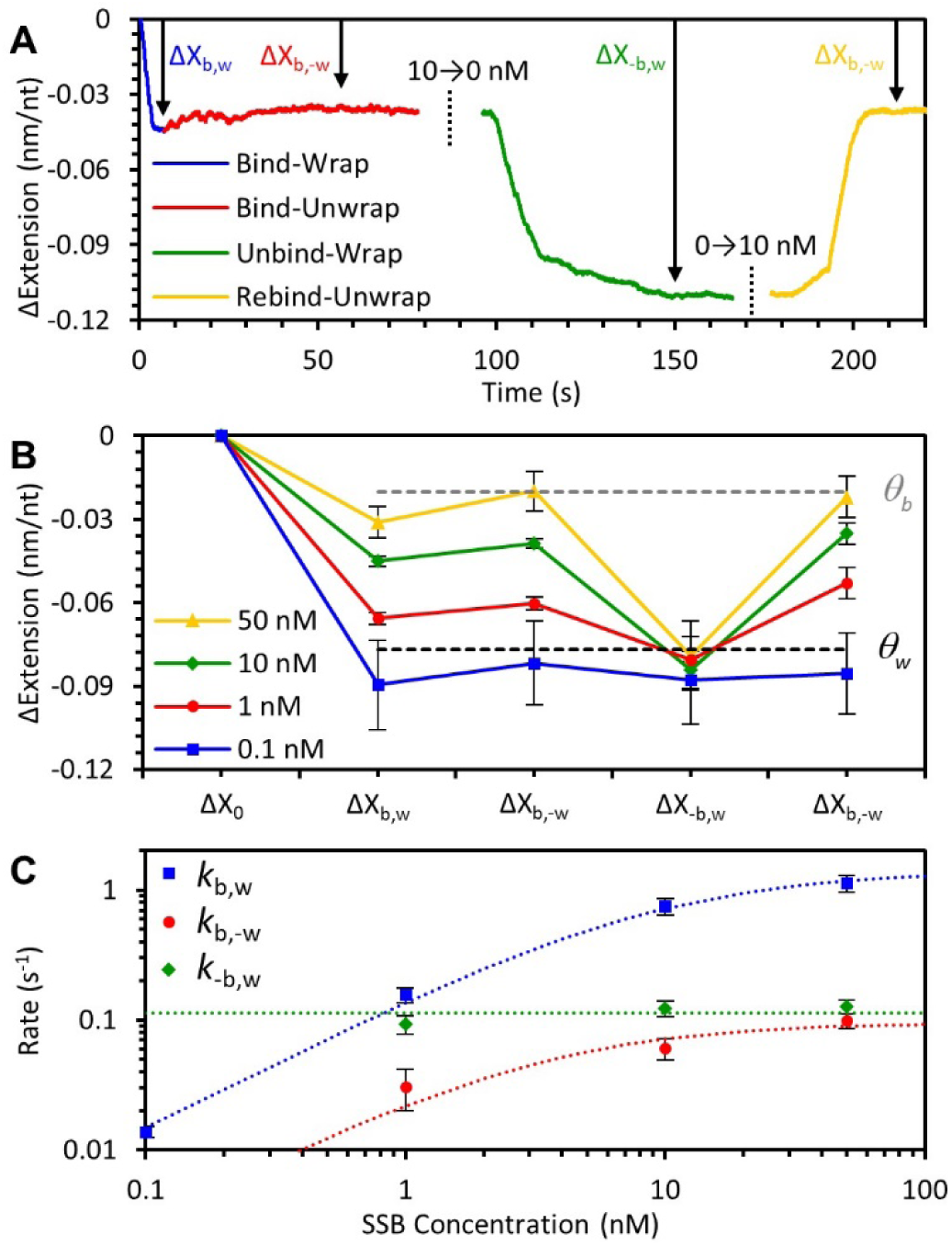
Concentration dependence of *Ec*SSB-ssDNA binding and wrapping at 12 pN template tension. (A) In the presence of free *Ec*SSB, the ssDNA extension first decreases due to the binding and subsequent wrapping of ssDNA by *Ec*SSB (bind-wrap,blue). A maximum extension decrease of ΔX_b,w_ is reached before the *Ec*SSB unwrapping outpaces wrapping due to continued binding (bind-unwrap,red), and the ssDNA extension change reaches a stable equilibrium of ΔX_b,-w_. Removal of free *Ec*SSB from solution results in some dissociation events without replacement (green). However, concomitant further wrapping of the bound *Ec*SSB results in a net extension-decrease of ΔX_-b,w_ (unbind-wrap). Reintroducing free protein allows *Ec*SSB to rebind to the *Ec*SSB-ssDNA complex that stimulates unwrapping events. This process (rebind-unwrap) registers as a net increase of ΔX_b,-w_ in the complex extension (magenta (B) The net change in extension of the *Ec*SSB-ssDNA complex after each step is averaged over multiple experiments for *Ec*SSB concentrations ranging from 0.1 to 50 nM. As the *Ec*SSB concentration is increased, the change in extension during incubation is decreased. Removal of free *Ec*SSB results in a consistent ∼0.08 nm/nt reduction in ssDNA extension (dotted line), regardless of initial *Ec*SSB concentration. The net extension change is consistent with the previously observed length change associated with a single *Ec*SSB_17_ on a 70 nt ssDNA substrate (27). Upon reintroducing free *Ec*SSB, the complex’s extension consistently reaches the same value as during the first incubation. (C) The average rate associated with each step of the *Ec*SSB-ssDNA interaction varies with free *Ec*SSB concentration. The bind-wrap step (blue) is rate limited by the initial binding of *Ec*SSB from solution at low concentrations, resulting in a linear dependence, and reaches an asymptote at high *Ec*SSB concentrations. The bind-unwrap step (red) is rate limited by the unwrapping events of *Ec*SSB, at a rate proportional to the bound but unwrapped *Ec*SSB fraction. The unbind-wrap step occurs at a constant rate of *k*_-b,w_=0.11 s^-1^ and is independent of the initial *Ec*SSB concentration (see Supplemental information for detailed derivation).

Since this bound but unwrapped *Ec*SSB state has been previously uncharacterized, we perform two additional experiments that prevent *Ec*SSB from wrapping (while leaving its ssDNA binding ability intact). First, we use a previously characterized *Ec*SSB mutant with the histidine at residue 55 replaced with tyrosine (H55Y) (31, 32), that does not form tetramers at low protein concentrations. Since the wrapped states of *Ec*SSB require ssDNA association with multiple OB domains of the tetramer, monomeric *Ec*SSB is unable to wrap ssDNA. Incubating the ssDNA held at 12 pN of tension with 5 nM monomeric H55Y mutant, yields a single-phase binding profile with a net extension change ∼0.015 nm/nt (Fig. S3A). This extension change is consistent with the equilibrium extension change that is observed with high concentrations of wild type (WT) *Ec*SSB (Fig. S3B). In the absence of free protein, H55Y dissociates from the ssDNA with a rate that is consistent with the direct dissociation observed with WT *Ec*SSB at higher forces (Fig. S3C). Moreover, the bimolecular on rate *k*_B_ of the monomeric H55Y at 5 nM agrees with the equivalent monomer concentration of the WT tetramer (1.25 nM) (Fig. 3C).

Second, we incubate *Ec*SSB with an ssDNA segment that is too short to wrap (<17 nt) and visualize these complexes using AFM imaging (Fig. S4). We incubated 100 bp dsDNA with an 8 nt poly dT overhang with *Ec*SSB at equimolar concentration (5 nM) in a buffer containing 10 mM Na^+^, 10 mM Mg^2+^, 10 mM Hepes, pH 7.5. The sample is deposited on an aminopropyltriethoxy silane (APS) coated mica surface. After imaging, both DNA constructs and *Ec*SSB tetramers were identified using a height threshold. *Ec*SSB binding specifically to the ssDNA can be readily identified by *Ec*SSB tetramers localizing at one end of the dsDNA region. This result confirms that *Ec*SSB can still bind ssDNA without wrapping.

### Binding and wrapping kinetics of *Ec*SSB

In order to measure the fundamental kinetic rates associated with *Ec*SSB dynamics, we fit the extension change over time for each phase of the binding experiment with a single-rate exponential function. These apparent rates are related to (but not exactly equal to) the fundamental rates of protein (un)binding and (un)wrapping, as defined by our model (see Supplemental Information). First, during the bind-wrap phase extension decreases as *Ec*SSB binds from solution then wraps the ssDNA. Thus, the measured rate (*k*_*b,w*_) is limited by both the rates of initial bimolecular binding (*c*·*k*_*b*_) and wrapping (*k*_*w*_). At low protein concentrations, *k*_*b,w*_ increases linearly with *Ec*SSB concentration *c*, yielding a bi-molecular rate of protein binding to bare ssDNA of *k*_*b*_*=*0.18 nM^-1.^s^-1^, while at higher protein concentrations it saturates at a constant value corresponding to the fundamental wrapping rate, *k*_*w*_=1.8 s^-1^ (blue line, Fig. 3C, see Supplemental Information for derivation). Second, during the bind-unwrap phase, ssDNA extension starts to increase as a consequence of unwrapping events of bound-*Ec*SSB, which allows further protein binding from the solution. Since this rate is nearly an order of magnitude slower than the rate of free protein binding (*k*_*b,-w*_≪*c*·*k*_*b*_), this process must be limited primarily by the rate of *Ec*SSB unwrapping, which asymptotes to a value of *k*_*-w*_=0.10 s^-1^ in the limit of high protein concentration (red line, Fig. 3C). Third, during the unbind-wrap phase the extension decreases as some *Ec*SSB dissociates, allowing other tetramers to wrap. Again, this process is much slower than the derived rate of wrapping (*k*_*-b,w*_≪*k*_*w*_), indicating this process is rate-limited by *Ec*SSB dissociation. Since this phase occurs in protein-free buffer, as expected this effective dissociation rate *k*_*-b*_=0.1 s^-1^ is independent of the initial *Ec*SSB concentration during incubation (green line, Fig. 3C). These fits yield values for the four fundamental rates that govern *Ec*SSB wrapping dynamics.

### Force-dependence of *Ec*SSB-ssDNA binding dynamics

Whereas we specifically detailed above *Ec*SSB wrapping dynamics while a force of 12 pN was maintained on the DNA substrate, these results are generalizable to other ssDNA tensions. We repeated the competitive binding measurements using 50 nM *Ec*SSB and observed biphasic binding at both lower (7 pN) and higher (20 pN) forces (Fig. 4A). The measured extension change increases with decreasing force, indicating that higher order wrapped states become progressively stable as the template tension is lowered. At 7 pN (blue line, Fig. 4B), the maximal extension change is observed after removing free protein to allow for maximal wrapping (ΔX_-b,w_=0.13 nm/nt). This extension change is consistent with the previously observed extension change associated with the *Ec*SSB_35_ mode as observed on a 70 nt ssDNA substrate (27) (Fig. S2). Therefore, at 7 pN, the maximally wrapped state in the absence of protein in solution corresponds to the *Ec*SSB_35_ proteins organized ∼70 nt apart on the ssDNA substrate. Nonetheless, the equilibrium complex extension change, ΔX_b,-w_ measured in the presence of 50 nM *Ec*SSB in solution at 7 pN is much smaller than the extension change expected for only the *Ec*SSB_17_ mode, indicating that both *Ec*SSB_8_ and *Ec*SSB_17_ states remain stable in the presence of these high concentrations of free protein. Thus, at 7 pN the *Ec*SSB-ssDNA complex primarily exists in the three alternative states: *Ec*SSB_8_, *Ec*SSB_17_, and *Ec*SSB_35_, where the relative occupancy of each state is governed by the concentration of free protein in the solution. While this three-state system description is more complicated, the kinetics and equilibrium properties of these complexes appear to be qualitatively similar to the ones we observe for two-state complexes at 12 pN as discussed above. At 20 pN (green line, Fig. 4B), wrapping is greatly destabilized, and most of the bound protein is unable to wrap, as evidenced by the minimal ssDNA compaction. Additionally, once free protein is removed, we measure a gradual extension increase over a ∼100 s timescale (Fig. 4C). The final extension approaches the extension of the protein-free ssDNA, indicating complete dissociation of *Ec*SSB. This is also supported by the observation that as the protein solution is re-introduced there is a biphasic binding profile (bind-wrap followed by bind-unwrap) that typically occurs during the initial protein incubation with protein-free ssDNA. Fitting an exponential rate to this process returns a much slower rate of dissociation *k*_*-b*_=0.017 s^-1^ with respect to the rate observed during the unbind-wrap transitions (0.1 s^-1^). Furthermore, we conducted force-jump experiments to test whether *Ec*SSB can remain bound to ssDNA at even higher tensions (Fig. S2). Here, first a maximally wrapped *Ec*SSB-ssDNA complex is produced at 12 pN by incubating ssDNA with 10 nM *Ec*SSB and then removing the free protein from the solution. The ssDNA is then abruptly (<1 s) stretched until a tension of 60 pN is obtained and held for 10 s, before bringing the tension back down to 12 pN. The ssDNA equilibrates to an extension slightly longer than that prior to the force-jump but remains significantly lower than that of of a protein-free ssDNA. Thus, while some *Ec*SSB dissociates during the force-jump, most remains bound and are able to rewrap when the ssDNA tension is brought back to 12 pN. This interpretation is further supported by a net increase in extension when protein is added back into the sample, indicating *Ec*SSB unwrapping events that are only observed on an *Ec*SSB-saturated ssDNA. We estimate the rate of protein dissociation (*k*_*-b*_) during the force jump by comparing the net ssDNA compaction due to wrapping just before and after the force-jump (*k*_*-b*_=0.017 s^-1^), which is consistent with both the directly observed rate of *Ec*SSB dissociation at 20 pN (Fig. S5), and with the monomeric H55Y at 12 pN (Fig. S3). Therefore, the direct dissociation rate, *k*_*-b*,_ is essentially force-independent. This result supports the hypothesis that in the *Ec*SSB_8_ mode only a single OB site on the *Ec*SSB tetramer is bound to ssDNA, resulting in minimal ssDNA deformation and compaction. In contrast, the various higher order wrapping modes in which *Ec*SSB greatly compacts ssDNA are therefore strongly destabilized by applied force on the ssDNA substrate.

**Figure 4:**
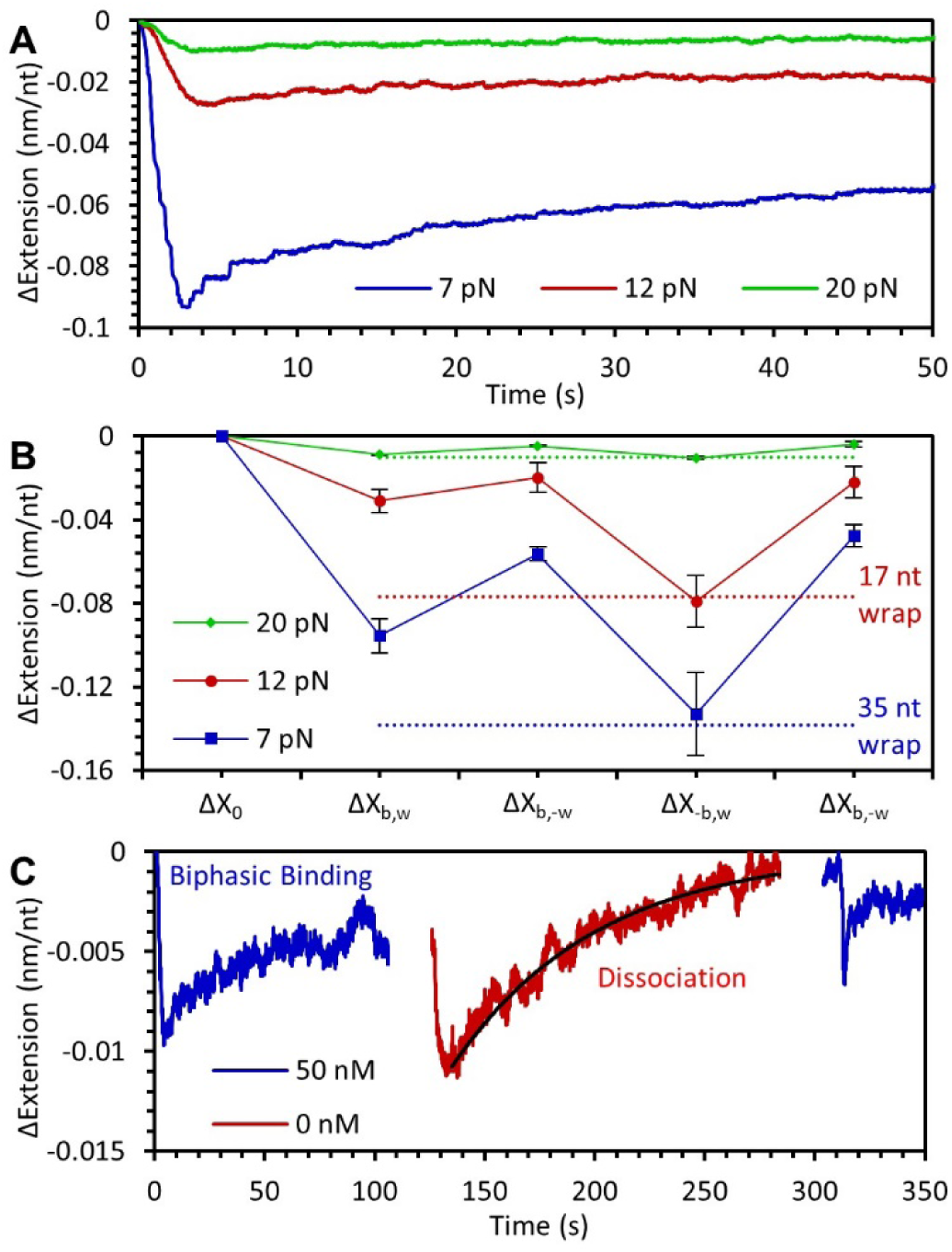
Force dependence of *Ec*SSB-ssDNA binding and wrapping. (A) As the force on the ssDNA template is decreased, the binding of 50 nM SSB causes a larger change in extension, consistent with *Ec*SSB accessing more wrapped states while bound to the ssDNA. A biphasic binding profile is seen at each force. (B) Average extension decrease at each phase of *Ec*SSB binding (compare to Fig. 3B) is shown for each force. After removing free protein (ΔX_-b,w_), the average extension decrease at 12 pN and 7 pN is consistent with an *Ec*SSB tetramer in the 17 nt wrapped state (red dotted line) and in the 35 nt wrapped state (blue dotted line), respectively, every 70 nt along the ssDNA substrate. (C) At 20 pN applied force, *Ec*SSB wrapping is unstable and most *Ec*SSB is in an unwrapped state. After removing free protein (red line), *Ec*SSB will dissociate from the ssDNA without replacement, leaving bare ssDNA. The ssDNA’s extension return to its original value is fit with an exponential (black line) to measure the rate of *Ec*SSB dissociation. Biphasic binding after the reintroduction of *Ec*SSB (second blue line), indicates the ssDNA is mostly free of protein after the dissociation step.

### Nearest neighbor interactions stimulate *Ec*SSB unwrapping and dissociation

In our various experiments, we measure two distinct rates of *Ec*SSB dissociation (Fig. 5A). The dissociation observed from an *Ec*SSB oversaturated complex, which is concomitant with further wrapping of the bound *Ec*SSB (achieved by oversaturating the ssDNA with high concentration *Ec*SSB, and then removing free protein), is faster, and occurs on the timescale of 10 s. In contrast, when wrapping is inhibited by high forces (F>15 pN) or when tetramerization (H55Y mutant) is inhibited, we observe much slower dissociation events (∼100 s) upon removing free protein from the solution. This former observation reflects the dissociation rate of *Ec*SSB in isolation, as the protein dissociation occurs from an unsaturated ssDNA substrate, with minimal nearest neighbor interactions. In contrast, the fast rate of dissociation is only observed when the ssDNA is oversaturated, when there are too many bound tetramers for the *Ec*SSB to wrap the ssDNA. Therefore, the dissociation of *Ec*SSB tetramers must be stimulated by the wrapping of the nearest neighbor competing proteins.

**Figure 5:**
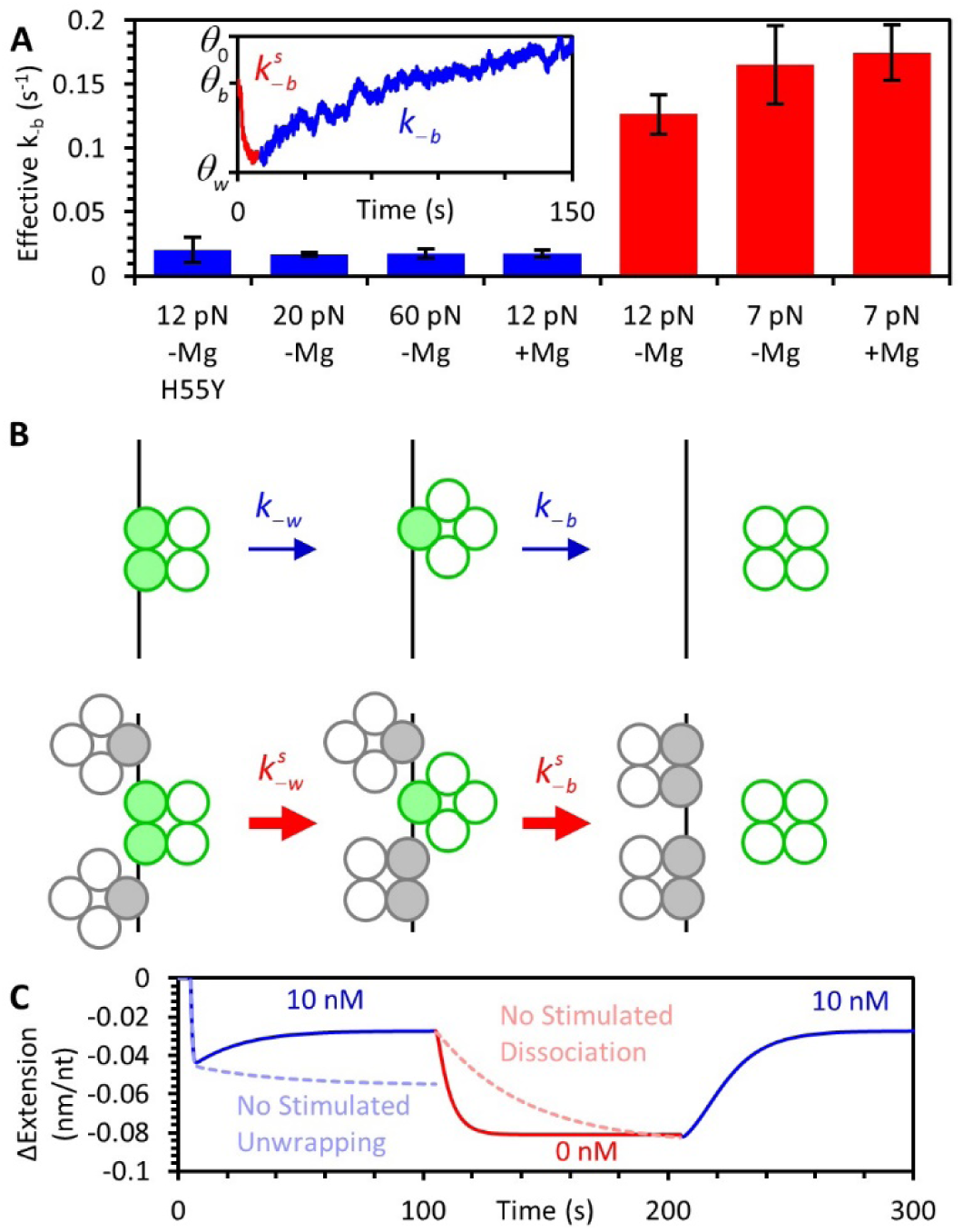
*Ec*SSB stimulated unwrapping and dissociation due to near neighbor effects. (A) Direct dissociation of *Ec*SSB is observed with H55Y due to compromised tetramer formation, with WT at forces >20 pN in the absence of Mg^2+^ or at forces >12 pN in the presence of 4 mM Mg^2+^. In either case the direct dissociation from unsaturated ssDNA complex is measured to be ∼0.015 s^-1^ (blue bars). In contrast, *Ec*SSB dissociation from an oversaturated ssDNA complex that is concomitant with wrapping events of neighboring *Ec*SSB occurs 10-fold faster at a rate of ∼0.15 s^-1^ (red bars). Inset shows the unbind-wrap process at 20 pN in which rapid stimulated dissociation (*k*^s^_-b_) from the oversaturated complex (red) is followed by the slow dissociation (*k*_-b_) events from the now unsaturated complex (blue)). (B) Schematic (ssDNA in black and *Ec*SSB in green) showing near neighbor stimulation of dissociation and unwrapping events. The net interacting interfaces of *Ec*SSB-ssDNA decreases upon an *Ec*SSB dissociation from an unsaturated ssDNA, leaving behind free ssDNA. However, in an oversaturated *Ec*SSB-ssDNA complex the net loss of protein-ssDNA interactions is minimal or none as near neighbors (gray circles) may compete to accommodate the substrate made available by unwrapping or dissociation events. (C) Model correction with simulated dissociation and unwrapping. The proposed two-step model (Fig. 2) reproduces the observed experimental *Ec*SSB-ssDNA wrapping kinetics (Fig. 1) in the presence and absence of free *Ec*SSB (blue and red lines) only when the stimulated dissociation and unwrapping is taken into account. Disregarding *k*_-b_ or *k*_-w_ stimulated dissociation and unwrapping (by keeping *k*_-b_ or *k*_-w_ constant) in the model results in a loss of the biphasic extension as seen with high *Ec*SSB concentrations (light blue dotted line), and a much slower unbind-wrap process (light red dotted line), which is inconsistent with the observed results (Figs. 1D,3A).

This fast rate of stimulated dissociation can be potentially explained by the energetic favorability of the various states of *Ec*SSB wrapping. When an isolated *Ec*SSB tetramer unwraps and dissociates from an ssDNA substrate, each associated OB domain must come off in succession until it is entirely free of the ssDNA (top path, Fig. 5B). This process, which results in a large region of bare ssDNA, is energetically unfavorable due to the high binding affinity of *Ec*SSB. In contrast, both unwrapping and dissociation are more favorable if the ssDNA is oversaturated with *Ec*SSB (bottom path, Fig. 5B). Under these conditions, both unwrapping and dissociation of one protein frees up ssDNA regions which can then be captured by neighboring proteins, maintaining an optimal number of individual domains bound to the ssDNA substrate.

We further show that both stimulated dissociation and stimulated unwrapping must be taken into account in order for the generalized two-step binding model to accurately reproduce our experimental data. Assuming *k*_*-b*_ and *k*_*-w*_ are constants and setting them to the values as determined in the absence of nearest neighbor interactions in the simulations eliminates the biphasic profile of the initial binding curve. Moreover, this assumption predicts a much slower unbind-wrap transition than what is observed (Fig. 5C). This is because, first, unwrapping is unable to outpace wrapping at high *Ec*SSB concentrations when the ssDNA is oversaturated, and second, further wrapping, observed when free protein is removed, requires a much longer timescale to re-equilibrate (Fig. 5C). Instead, the rates of both dissociation and unwrapping must effectively increase by an order of magnitude as the ssDNA substrate is oversaturated with *Ec*SSB.

### Progressively decreasing substrate triggers *Ec*SSB nearest neighbor interactions and dissociation

Our results demonstrate that an excess of free protein in solution leads to an oversaturated ssDNA substrate. Moreover, this oversaturation stimulates *Ec*SSB unwrapping events, favoring less wrapped *Ec*SSB states, in agreement with previous bulk solution observations (25). Alternatively to increasing free protein concentration, gradually decreasing the available length of ssDNA substrate, which occurs during replication, also increases the density of *Ec*SSB along the substrate. We achieved this effect by allowing irreversible RecA filamentation on a long *Ec*SSB-ssDNA complex. Because RecA-ssDNA filamentation requires Mg^2+^ cations, we first investigate *Ec*SSB-ssDNA binding dynamics in a solution containing Mg^2+^ (50 mM Na^+^, 4 mM Mg^2+^, 10 mM Hepes, pH 7.5). The overall binding dynamics of *Ec*SSB remain similar in the Mg^2+^ buffer (Fig. S6). In the presence of Mg^2+^ the local secondary structures slightly shortens the ssDNA molecule at lower forces (33) (Fig. S6A). Correcting for this additional extension change in the protein-free ssDNA extension yields the same equilibrium extension changes as observed in the Mg^2+^ free buffer (Fig. S6D). However, as the free protein in the solution is removed, the presence of Mg^2+^ resulted in *Ec*SSB dissociation at 12 pN in contrast to the Mg^2+^-free buffer, where we observe the unbind-wrap transition (Fig. S6B). Interestingly, the measured dissociation rate *k*_-b_ at 12 pN in the Mg^2+^ buffer is consistent with the same rate measured at 20 pN in the Mg^2+^ free buffer, suggesting that fluctuations between the bound *Ec*SSB_8_ and wrapped states are enhanced in the presence of Mg^2+^. We do not observe dissociation, however, at 7 pN, even in the presence of Mg^2+^. For this reason, we investigate the displacement of the ssDNA-bound *Ec*SSB by RecA filamentation at 7 pN of applied tension in the Mg^2+^ buffer (Fig. 6). First, we examine RecA filamentation on an *Ec*SSB-free ssDNA substrate. A Mg^2+^ buffer solution containing 100 nM RecA and 100 μM ATPγS is introduced to an ssDNA molecule held at 7 pN. The RecA-ssDNA nucleoprotein complex is formed via a slower nucleation step followed by a faster directional filamentation (34-37). As the filamentation proceeds, the increase in the flexibility of the RecA-ssDNA complex is registered as a gradual increase in the ssDNA extension that follows a single exponential behavior (Fig. 6A). Next, we repeat this experiment on an *Ec*SSB-ssDNA complex. To do so, we first incubate the ssDNA molecule that is held at 7 pN with 50 nM *Ec*SSB buffer in the Mg^2+^ buffer and subsequently rinse out the free *Ec*SSB from solution (Fig. 6B). After the unbind-wrap transition the *Ec*SSB-ssDNA complex stably equilibrates in its maximally wrapped state (predominantly with *Ec*SSB_35_ at 7 pN) for long timescales (∼1000 s) with no significant dissociation observed. Then we incubated the *Ec*SSB-ssDNA complex with a solution containing 100 nM RecA and 100 μM ATPγS to allow RecA filamentation, which proceeds at ∼10-fold slower rate than the filamentation observed on an *Ec*SSB-free ssDNA. The resulting protein-ssDNA complex after either procedure represents a completely RecA-filamented ssDNA, as evidenced by the subsequent force-extension cycle (38) (Fig. 6C). This indicates that RecA filamentation resulted in complete dissociation of *Ec*SSB that otherwise was highly stable in its maximally wrapped conformation. The total degree of RecA saturation can be calculated over the timescale of each experiment using the instantaneous ssDNA extension relative to the final extension (Fig. 6D). Fitting rates to these curves yields the rate of RecA filamentation both along protein-free ssDNA and *Ec*SSB-wrapped ssDNA (Fig. 6E). *Ec*SSB slows RecA filamentation by an order of magnitude as it becomes limited by the rate of *Ec*SSB dissociation. The progressing filamentation will gradually increase the *Ec*SSB:ssDNA ratio by reducing the ssDNA substrate that can be occupied by *Ec*SSB. The increased protein density will in turn trigger the stimulated unwrapping /dissociation events due to nearest neighbor interactions. If the progressing filamentation would have to wait for each *Ec*SSB protein to unbind with its stimulated dissociation rate of ∼0.1 s^-1^ at the site of filament growth, then the expected RecA filamentation rate would be (1/(8100 nt/70nt·10 s)) ∼0.0008 s^-1^, which is much slower than the observed ∼0.003 s^-1^ filamentation rate. This implies that as the protein density on the substrate increases the rate of *Ec*SSB dissociation along the whole ssDNA length maintains 70 nt per protein occupancy. The remaining *Ec*SSB slides along the ssDNA, releasing substrate for the RecA filament to proceed from a single or a few boundaries.

**Figure 6:**
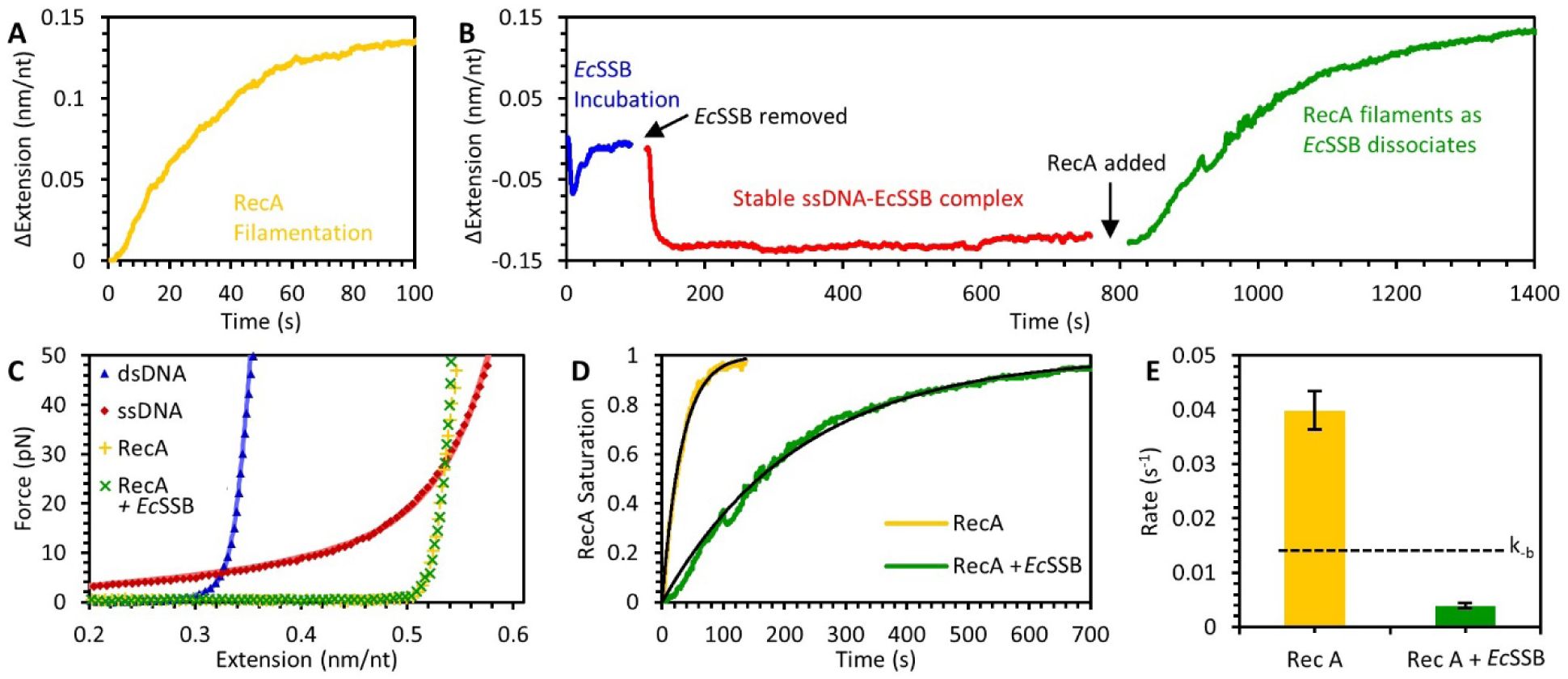
Dissociation of *Ec*SSB during RecA filament formation. (A) RecA filamentation (100 nM RecA with 100 µM ATPγS) on bare ssDNA at 7 pN occurs at a timescale of ∼10 s, and is registered as an increase in the ssDNA length due the increase in the its persistence length. (B) RecA filamentation on a maximally wrapped *Ec*SSB-ssDNA complex at 7 pN. The *Ec*SSB-ssDNA complex is obtained by first incubating the bare ssDNA at 7 pN with 50 nm *Ec*SSB and then removing the free *Ec*SSB from the solution as described in the text. Here, RecA forms filaments, presumably displacing EcSSB from ssDNA, but at a much longer (∼100 s) timescale. (C) The resultant force-extension profiles of the RecA-ssDNA filaments formed in both (A) and (B) are identical, which confirms complete RecA filamentation in either case. (D) Normalized extension-time profiles of RecA filamentation kinetics on the bare ssDNA (yellow) and *Ec*SSB-ssDNA complex (green) yield simple exponential functions (black) (E) Comparison of the RecA filamentation rates in (A) and (B) shows that the RecA filamentation on the *Ec*SSB-ssDNA complex is rate limited by the *Ec*SSB dissociation rate, *k*_-b_ (dashed black line).

## Discussion

### *A* competitive binding mechanism allows oversaturation and stimulates dissociation

We show that the ssDNA is oversaturated via stimulated unwrapping when the free protein is abundant in solution. However, through simulated dissociation, this complex equilibrates to a configuration that accommodates one wrapped *Ec*SSB per 70 nt of ssDNA substrate when excess free protein is removed in a non-cooperative manner. Furthermore, we find that the protein concentration leading to half displacement of the *Ec*SSB_17_ with *Ec*SSB_8_ complexes (*c\**∼3 nM, Fig. 2C, see Supplemental Infomration) is significantly higher than the equilibrium dissociation constant that we measure for *Ec*SSB_8_ binding to our long ssDNA without free ends (*K*_*d*_ = 0.1 nM). This observation strongly supports the idea that the weaker *Ec*SSB_8_ binding affinity and its faster dissociation from the oversaturated *Ec*SSB-ssDNA complex is a consequence of competitive displacement of ssDNA from the less-wrapped *Ec*SSB by its nearest-neighbor *Ec*SSB. Using RecA filamentation, we show that increasing the *Ec*SSB density on the ssDNA substrate results in complete dissociation of *Ec*SSB that is otherwise stable for ∼100 s. This dissociation occurs over the entire length of the ssDNA in an anti-cooperative manner, as discussed in the next section. This contrasts with what is expected from a cooperatively bound protein, for which its higher level of occupancy is expected to lead to slower dissociation due to the fewer boundaries between bound and unbound domains. Therefore, we suggest that *Ec*SSB binding dynamics is not cooperative, but instead represents a competitive process that allows rapid kinetics during genomic maintenance, as discussed in the next sections.

### Stimulated unwrapping and dissociation are near isoenergetic processes

We show that the kinetics and equilibrium of *Ec*SSB binding to “unsaturated” vs “oversaturated” ssDNA differ dramatically, resulting in stimulated dissociation and stimulated unwrapping events only from the oversaturated state. The oversaturated complex occurs when more than one *Ec*SSB tetramer is bound per ∼70 nt (Figs. S1 and S2). Thus we hypothesize that the two ssDNA-bound *Ec*SSB tetramers are no longer influenced by each other once the average distance separating them becomes larger than the typical length (*L*) that the *Ec*SSB can diffuse on ssDNA during the time ∼10 s of its stimulated dissociation, which can be estimated to be (*L ≥ D*·*t)*^*0.5*^ ∼ (300 nt^2^/s·10 s)^0.5^)∼ 55 nt, where D is the *Ec*SSB diffusion coefficient on ssDNA (30). While this is a rough estimate, it yields a plausible explanation for the *Ec*SSB density on ssDNA that separates its unsaturated and oversaturated binding regimes.

Interestingly, we find that the stimulated dissociation rate, *k*^*s*^_*-b*_ is close in magnitude to the *Ec*SSB_17_ stimulated unwrapping rate, *k*^*s*^_*-w*_ ∼ 0.10 s^-1^, suggesting that these two rates might be limited by the timescale for ssDNA to peel from a single *Ec*SSB OB site. Simultaneous rebinding of this released ssDNA by another neighboring *Ec*SSB may stimulate the dissociation of an *Ec*SSB_8_. This is an almost isoenergetic process that does not require complete OB-ssDNA dissociation, and is, therefore, expected to be much faster. We suggest that the mechanism of stimulated dissociation from its oversaturated ssDNA complex is similar to the previously established mechanism of rapid diffusion of tightly wound *Ec*SSB on bare ssDNA (30). *Ec*SSB diffusion on ssDNA was shown to proceed via a reptational mechanism without protein unwinding or dissociation. This process was shown to be much faster than the ssDNA wrapping dynamics, as it does not involve releasing all the OB sites from ssDNA. Instead these bonds are partially released (2-5 nts), and immediately replaced by adjacent nts. This process is near isoenergetic and results in much faster *Ec*SSB motion on ssDNA, while remaining fully bound and wrapped. Similarly, the stimulated *Ec*SSB dissociation and unwrapping on an oversaturated ssDNA complex is much faster than from an unsaturated ssDNA, as the *Ec*SSB-ssDNA bonds on one *Ec*SSB are being gradually replaced by similar interactions with its nearest neighbor.

### Rapid *Ec*SSB kinetics ensures maximum ssDNA coverage during genomic maintenance

The ssDNA exchange rate is regulated by the ongoing coordinated enzymatic activity of the DNA polymerases, helicases, RecA, etc. during genomic maintenance processes. Despite extensive studies of *Ec*SSB with both traditional biochemical approaches in bulk solutions, as well as with single molecule approaches on short stretched ssDNA accommodating one *Ec*SSB, the following fundamental question still remains unanswered. How do *Ec*SSB tetramers that are tightly bound and highly wound on ssDNA undergo the rapid complex rearrangements involving protein dissociation and re-association that are required to keep up with the enzymes involved in genomic maintenance? The current study provides critical new information to help answer this question.

As the amount of *Ec*SSB in bacteria is kept at the level sufficient for saturation of all available ssDNA (39-41), there should be a constant exchange of the ssDNA substrate within the saturated complex with *Ec*SSB. As most ssDNA is always *Ec*SSB-saturated, the mechanism of rapid *Ec*SSB diffusion on bare ssDNA (30) likely does not contribute significantly to the rapid *Ec*SSB turnover. Also, a massive *Ec*SSB transfer over long distances between the saturated and bare ssDNA is unlikely to be the main mechanism of such *Ec*SSB turnover, as it requires these distant ssDNA regions to be in direct contact with each other. Our results suggest novel pathways of the efficient *Ec*SSB exchange between distant ssDNA regions, that involves fast *Ec*SSB dissociation into bulk solution followed by the fast re-association with either saturated or bare ssDNA. If ssDNA is available nearby for rebinding, this would allow *Ec*SSB recycling, as suggested by a recent single molecule study (42).

Taken together, the results described in this study provide mechanisms to regulate the density of the ssDNA-bound *Ec*SSB, which is central for its transient role during genome maintenance and replication. Based on the findings in this study, we propose a mechanism of self-regulation of protein density, a phenomenon that emerges directly from competitive *E*cSSB binding dynamics (Fig. 7). Here we consider the case of genomic replication with the assumption that the template ssDNA is saturated with *Ec*SSB in an intermediately wrapped state such as *Ec*SSB_35_, as was previously seen with the human mitochondrial SSB that is structurally and functionally similar to *Ec*SSB (43). However, we must also note that *Ec*SSB exists in a dynamic equillibrium between its distinct modes that are able to diffuse along the DNA without dissociation (29, 43). Therefore, a processing enzyme, the DNA polymerase (DNA pol) in this case, may displace the wrapped *Ec*SSB during synthesis, pushing it forward along the template strand. Supporting this idea, a recent study investigated the species-specific inter-protein interactions between the DNA pol and SSB that enhance the SSB displacement by DNA pol (44). The active displacement of *Ec*SSB by the DNA pol along with *Ec*SSB’s ability to slide along the ssDNA (30) may increase the *Ec*SSB density throughout the template strand. This may in turn produce an instantaneous oversaturated *Ec*SSB-ssDNA complex, in which the stimulated unwrapping events will be followed by stimulated dissociation events along the entire template strand. As the ssDNA template gets smaller, this dynamic process may continue and self-regulate the *Ec*SSB density efficiently to allow the DNA pol to proceed while ensuring maximal template coverage at any given time. The proposed mechanism is entirely based on the *Ec*SSB’s competitive binding mechanism to ssDNA, and its nearest neighbor interactions that allow oversaturation while stimulating dissociation. Future studies may provide insights on the structure-function relationship and the molecular mechanism by which these nearest neighbor interactions are mediated.

**Figure 7:**
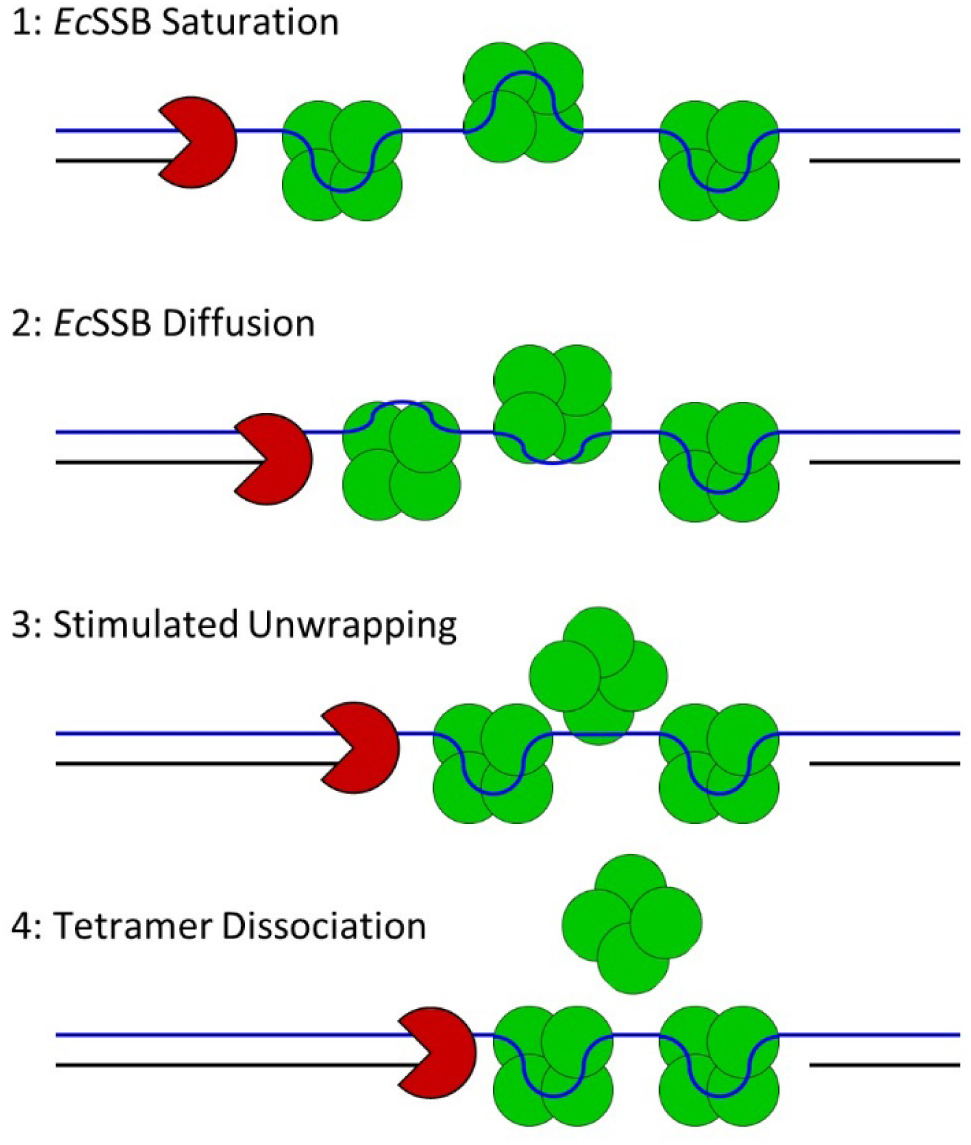
Self-regulation of protein density mechanism. (1) A transiently formed ssDNA substrate is saturated with one *Ec*SSB per 70 nt. (2) A processing enzyme such as the DNA polymerase will gradually decrease ssDNA substrate, and due to diffusion of an oversaturated *Ec*SSB-ssDNA complex. (3) As the instantaneous *Ec*SSB density is increased resulting in more than one *Ec*SSB per 70 nt, near neighbor effects stimulate the unwrapping events. (4) Stimulated dissociation concomitant with near neighbor wrapping events removes excess *Ec*SSB from the ssDNA substrate. This process can happen along the entire *Ec*SSB-ssDNA complex length, simultaneously dissociating many EcSSB tetramers and thereby self-regulating its density. This mechanism ensures fast turnover of ssDNA substrate to the processing enzyme as it translocates along ssDNA, while providing maximal ssDNA coverage at any given time

## Materials and Methods

### Preparation of DNA substrates and proteins

For the optical tweezer experiments, an 8.1 kbp dsDNA construct with digoxigenin (DIG) and biotin labeled ends with a free 3′ end was constructed as previously described (45). Vector pBACgus11 (gift from Borja Ibarra) was linearized through double digestion using restriction enzymes SacI and BamHI (New England Biolabs, NEB). A dsDNA handle with digoxigenin (DIG) labeled bases with a complementary end to the BamHI sequence was PCR amplified (46). The DIG handle and a biotinylated oligo (Integrated DNA Technologies, IDT) were annealed to the overhangs produced by BamHI and SacI then ligated using T4 DNA ligase (NEB).

For the AFM experiments, a hybrid dsDNA-ssDNA construct was produced, which enables accurate detection of protein binding to an ssDNA substrate (47). A PCR amplified dsDNA segment from pUC19 was digested by BamHI and ligated to an oligo with a complementary end (IDT) using T4 DNA ligase. The final product consisted of 100 bp of dsDNA with an 8 nt long poly dT tail.

WT *Ec*SSB and T7 DNA polymerase were purchased (NEB). The plasmid encoding WT *Ec*SSB pEAW134 was a gift from Dr. Mark Sutton of the University at Buffalo. The *Ec*SSB_H55Y_ variant was constructed using Quikchange site-directed mutagenesis (Agilent) and mutagenic oligonucleotides. Recombinant proteins EcSSB_WT_ and EcSSB_H55Y_ were expressed in *E. coli* BL21 Tuner cells in 1 L Luria Broth with ampicillin (100 µg/mL). After the cells reached an OD_600_ of ∼0.7 expression was induced by adding IPTG to a final concentration of 1 mM and shaking at 220 rpm for 4 h at 30 °C. Purification was carried out based on the protocols outlined by Lohman et. al. with some modification (48). All subsequent steps were carried out at 4 °C or on ice. For EcSSB_WT_ cells were collected by centrifugation and resuspended in 20 mL of buffer containing 50 mM Tris pH 8.3, 200 mM NaCl, 15 mM Spermidine trichloride, 1 mM EDTA, 100 µM PMSF and 10% sucrose. Lysis was carried out via sonication and the addition of lysozyme. Cells containing EcSSB_H55Y_ were handled similarly except for an increase in salt to 400 mM NaCl to induce the alternate DNA binding mode (SSB)_65_ to compensate for the reduced binding affinity of the H55Y variant (24). The collected supernatant was subjected to Polymin P (Sigma Aldrich) precipitation by adding a 5% solution dropwise to a final concentration of 0.4%. Stirring was continued for 20 min before centrifugation at 10,000 xg for 20 min. The resulting pellet was collected and resuspended gently in 50 mM Tris pH 8.3, 400 mM NaCl, 1 mM EDTA, and 20% glycerol to the initial fraction volume over 60 min followed by centrifugation at 10,000 xg for 20 min. SSB was precipitated from the collected supernatant by slowly adding ammonium sulfate (Sigma Aldrich)with stirring to a final concentration of 150 g/L and manually stirring for an additional 30 min followed by centrifugation at 24,000 xg for 30 min. The resulting pellet was resuspended in 50 mM Tris pH 8.3, 300 mM NaCl, 1 mM EDTA, and 20% glycerol at 0.9x fraction volume. Purity was examined by SDS-PAGE and concentration determined by Bradford assay before loading onto a 20 mL spin column packed with 5 mL ssDNA-cellulose (Sigma Aldrich D8273-5G). The column with SSB containing fractions was sealed and incubated for 60 min with gentle rocking. Washing and elution were carried out by centrifugation at 1000 xg and the duration of each centrifugation event was determined prior to loading the protein in order to prevent drying the column. The buffer used for wash and elution steps was 50 mM Tris pH 8.3, 1 mM EDTA, 20% glycerol, and NaCl at 300 mM, 600 mM or 2 M. After allowing the column to drain it was washed with 10 CV of 300 mM NaCl buffer, then 10 CV of 600 mM NaCl buffer followed by elution with 10 CV of 2 M NaCl. Fractions were evaluated by SDS-PAGE and the 2 M NaCl elution fractions containing SSB were pooled together before determining concentration by Bradford assay and concentrating by ammonium sulfate precipitation at 225 g/L. The resulting pellet is then resuspended in 50 mM Tris pH 8.3, 300 mM NaCl, 1 mM EDTA, 1 mM β-Mercaptoethanol and 50% glycerol to the desired concentration.

#### Optical tweezers

The 8.1 kbp dsDNA construct with a primer-template junction at one terminus was tethered between 2 μm anti-digoxigenin and 3 μm streptavidin functionalized beads (Spherotech) held in place by a micropipette tip and a dual beam optical trap, respectively. The micropipette tip was moved by a piezo electric stage with 0.1 nm precision to change the extended length of the DNA while the deflection of the laser trap was measured to calculate the force exerted on the trapped bead and thus the tension along the DNA. Additionally, a bright-field image of the two beads was recorded at 40X magnification. The instrument is controlled via a NI-DAQ interface (National Instruments) and custom software compiled with LabWindows. In order to create an ssDNA binding template, T7 DNA polymerase (T7DNAp) was introduced into the sample and the DNA was held at a constant force of 50 pN to trigger exonucleolysis (49) and completely digest one strand to produce a long ssDNA. After thorough rinsing of the T7DNAp reaction buffer DNA was held at fixed forces in a buffer containing 10 mM HEPES, 50 mM Na+, pH 7.5, except where specifically noted. Then *Ec*SSB was introduced to the cell and the position of the bead was continuously adjusted via a force feedback loop to maintain constant DNA tension. Free *Ec*SSB in the solution was removed by replacing the protein solution with a protein free buffer. After data acquisition, the relative distance between the beads was calculated using the bright-field images and compared to the extension of the DNA as calculated by the position of the piezo electric state. This comparison allows to correct for long term thermal drifts of the cell flow cell system. All the data were analyzed using custom code written in MATLAB (Mathworks).

#### AFM Imaging

*Ec*SSB and dsDNA-ssDNA hybrid constructs were incubated at an equimolar ratio (5 nM) in a buffer containing 10 mM Na^+^, 10 mM Mg^2+^, and 10 mM Hepes, pH 7.5. The sample was deposited on an APTES coated functionalized mica surface (50) and then imaged in fluid using peak force tapping mode (Bruker). Images were analyzed using Gwyddion software and height thresholds of 0.5 and 1.5 nm were used to identify dsDNA markers and EcSSB tetramers respectively.

## Acknowledgements

We thank Michelle Silva and students in the Chemical Biology class for constructing the SSB H55Y variant.

