## Supplement for "Self-regulation of single-stranded DNA wrapping dynamics by *E. coli* SSB promotes both stable binding and rapid dissociation"

### *Supplemental Information*

**Supplement 1. Explicit analytical description of the kinetics and equilibrium of *Ec*SSB binding and competition between its different states on ssDNA.**

***Kinetics of EcSSB-ssDNA complexes during the protein concentration jump cycles.***

The bind-wrap transition followed by the unwrap-bind transition is a complicated process with no explicit analytical solution. However, our understanding of the underlying elementary processes allows us to express the main component of each observed rate through the elementary reaction rates of this system. As discussed in the main text, at 12 pN these transitions can be primarily denoted with the following kinetic scheme.

$$EcSSB_{unbound} \underset{k_{-b}}{\overset{ck_b}{\rightleftharpoons}} EcSSB_8 \underset{k_{-w}}{\overset{k_w}{\rightleftharpoons}} EcSSB_{17}. \qquad \text{S1}$$

Here, $ck_b$ and $k_{-b}$ are the forward and reverse rates of free protein binding and $k_w$ and $k_{-w}$ are the forward and reverse rates of bound protein wrapping the ssDNA substrate. Experimentally, we investigate this system by varying the free protein concentration, *c*, that presumably mimics the variations in *Ec*SSB concentration in the bacterial cell due to the fluctuations in protein expression and the availability of transient ssDNA during genomic maintenance. For the sake of simplicity, we derive the analytical expressions describing the kinetics and equilibrium of the three-state system as observed at a template tension of 12 pN where primarily *Ec*$SSB_8$ and *Ec*$SSB_{17}$ states coexist. At lower forces the coexistence of several higher order wrapped *Ec*SSB states would make deconvolving the system into its fundamental reaction steps mathematically challenging. However, the kinetic scheme described here is qualitatively consistent at lower template tensions where more than two *Ec*SSB states coexist. For example, at a template tension of 7pN we observe the same qualitative behavior (Fig. 4), but the model would have to be expanded to incorporate higher order states such as *Ec*$SSB_{35}$, and *Ec*$SSB_{56}$ in addition to the *Ec*$SSB_8$ and *Ec*$SSB_{17}$ states (Fig. S2). As described in the text we find that the *Ec*SSB dissociation and unwrapping kinetics depends on the degree of ssDNA saturation. We show that the *Ec*SSB-ssDNA complex stably equilibrate with one *Ec*SSB per ~70 nt (Fig. S1, S2). Thus, when there are more/less *Ec*SSB per ~70 nt the complex is defined to be *Ec*SSB oversaturated/unsaturated. Thus, the net dissociation and unwrapping rates are given by,

$$k_{-b} = k^0_{-b} + k^s_{-b} \quad and \quad k_{-w} = k^0_{-w} + k^s_{-w}. \qquad \text{S2}$$

Here, the superscripts 0 and s represent an *Ec*SSB -unsaturated and -oversaturated complex, respectively.

The explicit expressions for the observed bind-wrap ($k_{b,w}$), bind-unwrap ($k_{b,-w}$), and unbind-wrap ($k_{-b,w}$) rates in terms of the four fundamental forward and reverse kinetic rates are derived as a function of the free *Ec*SSB concentration in solution. During the bind-wrap transition *Ec*SSB binds and subsequently wraps an unsaturated ssDNA in series, and therefore the observed rate, $k_{b,w}$ can be estimated by,

$$\frac{1}{k_{bw}}=\frac{1}{ck_b}+\frac{1}{k_w} \quad \rightarrow \quad k_{bw}(c)=\frac{ck_bk_w}{ck_b+k_w}=\frac{k_w}{1+\frac{k_w}{ck_b}}. \qquad \text{S3}$$

Fitting the observed bind-wrap transition (Fig. 3C blue) to Eq. S3 yields, $k_b$ = 0.18 nM$^{-1}\cdot$s$^{-1}$ and $k_w$ ~ 1.3 s$^{-1}$

The subsequent bind-unwrap transition occurs as the ssDNA substrate becomes *Ec*SSB oversaturated. Thus, the observed rate $k_{b,-w}$ for this transition is given by,

$$\frac{1}{k_{b,-w}}=\frac{1}{ck_b}+\frac{1}{k_{-w}} \quad \rightarrow \quad \frac{ck_bk_{-w}}{ck_b+k_{-w}}=\frac{k_{-w}}{1+\frac{k_{-w}}{ck_b}}\approx\frac{k^s_{-w}}{1+\frac{k^s_{-w}}{ck_b}} \qquad \text{S4}$$

Experimentally, we do not observe unwrapping on an unsaturated ssDNA substrate at 12 pN as evidenced by the maximally wrapped *Ec*SSB-ssDNA complex being stable >100 s upon the removal of free *Ec*SSB in the solution (Fig. 3A, green line). We observe the unwrapping transition due to nearest neighbor interactions at sufficiently high protein concentrations when the ssDNA substrate is oversaturated (Fig. 1C). For this reason, the final approximation in Eq.S4 holds because the unwrapping rate on a saturated ssDNA substrate (stimulated unwrapping) must be at least an order of magnitude faster than that of on an unsaturated ssDNA substrate. Accordingly, at saturation stimulated unwrapping becomes the rate-limiting step and the fit yields, $k^s_{-w}$ ~ 0.10 s$^{-1}$ (Fig. 3C red).

Finally, the rate of the unbind-wrap transition in which an *Ec*SSB$_8$ dissociation is followed by the further wrapping of a neighboring protein is given by,

$$\frac{1}{k_{-bw}}=\frac{1}{k^s_{-b}}+\frac{1}{k_w} \quad \rightarrow \quad k_{-bw}(c)=\frac{k^s_{-b}}{1+\frac{k^s_{-b}}{k_w}}\approx k^s_{-b}. \qquad \text{S5}$$

As the unbind-wrap transition occurs on an *Ec*SSB-oversaturated ssDNA substrate, $k^s_{-b}$ represents the stimulated dissociation rate due to nearest neighbor interactions. Therefore, here we measure the rate of *Ec*SSB dissociation from an oversaturated ssDNA substrate to be $k^s_{-b}$=0.12 s$^{-1}$ (Fig. 3C green). Note that we directly measure the force-independant dissociation rate of *Ec*SSB from an unsaturated ssDNA to be $k^0_{-b}$=0.014 s$^{-1}$ (Figs.4C and S5). Therefore, the stimulated unwrapping and stimulated dissociation on a saturated ssDNA substrate are at least an order of magnitude faster than the equivalent processes on an unsaturated substrate.

***Competition of different EcSSB states within the equilibrated oversaturated EcSSB-ssDNA complex.***

From the association and dissociation rates observed on an unsaturated ssDNA substrate we can determine the equilibrium dissociation constant, $K_{db}$ of *Ec*SSB binding to an *Ec*SSB$_8$ state to be,

$$K_{db} = \frac{k_{-b}^{0}}{k_b} = \frac{0.014\, s^{-1}}{0.18\, nM^{-1} s^{-1}} = 0.08\, nM, \qquad \text{S6}$$

and the corresponding free energy to be,

$$G_{b,0}(k_B T) = \ln\left(\frac{c}{K_{db}}\right). \qquad \text{S7}$$

In the concentration-jump experiments (Fig. 3) the *Ec*SSB-ssDNA complex remains oversaturated between the bind-wrap transition through the quasi-equilibrium bind-unwrap transition up until the maximum winding after dissociation during the unbind-wrap transition. ($\Delta X_{b,w} \rightarrow \Delta X_{b,-w} \rightarrow \Delta X_{-b,w}$ in Fig. 3A-B). The equilibrium complex extension upon the bind-unwrap transtion, $\Delta X_{b,-w}$ (Fig. 3A-B) is defined by the fractions, [illegible] of saturated ssDNA engaged with either of the *Ec*$SSB_8$ ($\Theta_b$), or *Ec*$SSB_{17}$ ($\Theta_w$) states where,

$$\Delta X = -[\Delta x_b \Theta_b + \Delta x_w \Theta_w]. \qquad \text{S8}$$

Here [illegible]*x* represents the corresponding ssDNA length-change associated with each binding state. We can express these equilibrium competing fractions of *Ec*SSB states analytically in terms of their binding free energies and fundamental reaction rates as a function of free *Ec*SSB concentration (c) at 12 pN. We assume that in equilibrium at 12 pN the ssDNA substrate is bound by either *Ec*$SSB_8$ (B), or *Ec*$SSB_{17}$ (W) *Ec*SSB states. Even though the free energy of W state per protein, $G_b + G_w$ is larger than the free energy of B state per protein $G_b$, a single *Ec*SSB in W can be substituted by multiple B complexes with the smaller binding free energy due to differences different binding site sizes (N), where $N_w/N_b = 1/\eta > 1$. The equilibrium B and W fractions of associated with ssDNA substrate per $N_w$ nt is given by,

$$\eta.\overline{B} + \overline{W} = 1 \qquad \text{S9}$$

$\Theta_w$ and $\Theta_b$ are the ssDNA fractions engaged with the W and B states, respectively. Therefore,

$$\Theta_w = \overline{W} = \frac{e^{(G_b+G_w)/k_B T}}{Z} = \frac{1}{1+e^{-\delta G/k_B T}}, \qquad \text{S10}$$

and

$$\Theta_b = \eta.\overline{B} = \frac{e^{(G_b/\eta)/k_B T}}{Z} = \frac{1}{1+e^{\delta G/k_B T}}, \qquad \text{S11}$$

where

$$Z = e^{(G_b+G_w)/k_B T} + e^{(G_b/\eta)/k_B T} \quad and \quad \delta G = G_w - G_b\left(1/\eta - 1\right). \qquad \text{S12}$$

$\delta G$ is the free energy difference per $N_w$ of ssDNA length between W- and B- saturated states. The midpoint of the transition with equal fractions of $\eta B$=$W$=1/2 occurs when

$$\delta G = 0 \quad or \quad G_w = G_b\left(1/\eta - 1\right) \qquad \text{S13}$$

The later condition defines the midpoint free *Ec*SSB concentration, *c*(F)* at which *Ec*$SSB_8$ and *Ec*$SSB_{17}$ states are equally probable where,

$$c^* = \frac{k_{-b}}{k_b}.\left(\frac{k_w}{k_{-w}^s}\right)^{\eta/1-1} = K_{db}\left(\frac{k_w}{k_{-w}^s}\right)^{\eta/1-1} \qquad \text{S14}$$

The equilibrium fractions of ssDNA associated with the W and B state as given by Eq. S10 and Eq. 11 can be now written as,

$$\Theta_w = \overline{W} = \frac{1}{1+\frac{k_{-w}^s}{k_w}.\left(\frac{K_{db}}{c}\right)^{1/\eta-1}} = \frac{1}{1+\left(\frac{c}{c^*}\right)^{1/\eta-1}} \qquad \text{S15}$$

$$\Theta_b = \eta\overline{B} = \frac{1}{1+\frac{k_w^s}{k_{-w}}.\left(\frac{K_{db}}{c}\right)^{1/\eta-1}} = \frac{1}{1+\left(\frac{c^*}{c}\right)^{1/\eta-1}} \qquad \text{S16}$$

We model the observed $\Delta X_{b,-w}$ ($\equiv \Theta_b$) substituting the experimentally determined, $k_w$ (=1.3 $s^{-1}$), and $k^s_{-w}$ (=0.1 $s^{-1}$) by letting $\eta$ and $K_{db}$ be free parameters (Fig. 2C, solid line). The fit yields ~ 0.6, which is consistent with $N_w$ and $N_b$ being ~17 nt and ~8 nt, respectively. The fit yields $K_{db}$~ 0.04 nM, which is comparable to the measured $K_{db}$ (= 0.08 nM, Eq. S6). This suggests that the on/off kinetics of *Ec*SSB binding to unsaturated ssDNA is consistent with the competitive titration data in Fig. 3C. The transition midpoint concentration, *c** at which the bound *Ec*$SSB_8$ displaces the wound *Ec*$SSB_{17}$ can be estimated to be, ~3 nM. This value of c* is much higher than the actual $K_{db}$ (~ 0.08 nM) of *Ec*$SSB_8$ due to the competition with wrapped *Ec*SSB states. The other manifestations of the same competition are the much faster stimulated dissociation and stimulated unwrapping events on an oversaturated ssDNA, as discussed in the main text.

**Supplement 2. Numerical solutions of the two-step competitive binding model.**

To test the proposed three-state kinetic model, we first derive differential equations based on the kinetic scheme described in Eq. 1, which determines the rate of change in the fraction of ssDNA substrate that's is in the unbound ($\Theta_0$), bound ($\Theta_b$), and wrapped ($\Theta_w$) states as a function of the fundamental rates of *Ec*SSB binding, dissociation, wrapping, and unwrapping.

$$\frac{d\Theta_0}{dt} = \Theta_b k_{-b} - \Theta_0 c k_b + \Theta_w k_{-w}(1-\eta) - \Theta_0 \Theta_b k_w (\eta^{-1} - 1)$$
$$\frac{d\Theta_b}{dt} = \Theta_0 c k_b - \Theta_b k_{-b} + \Theta_w \eta k_{-w} - \Theta_0 \Theta_b k_w \quad \text{S17}$$
$$\frac{d\Theta_w}{dt} = \Theta_0 \Theta_b \eta^{-1} k_w - \Theta_w k_{-w}$$

Note that we are specifically modelling *Ec*SSB binding dynamics at 12 pN, where *Ec*$SSB_{17}$ is the only stably wrapped state. Modeling higher order wrapped states would require adding additional terms for the occupancy of these states. We substitute fundamental rates as determined by our concentration dependent binding experiments (Fig. 3) and solve the differential equations using the Euler method for small discrete time steps (1 ms). This yields the fractions of ssDNA in each state over time for a given free concentration of protein. Additionally, the free protein concentration is abruptly changed to zero after the incubation step is complete to simulate the concentration-jump experiments. Finally, the occupancy of each state over time is converted into an observable change in ssDNA extension for direct comparison with experimental data. The total extension change as a function of time due to *Ec*SSB binding, ΔX (t), is given by

$$\Delta X(t) = -\left[\Delta X_b \Theta_b(t) + \Delta X_w \Theta_w(t)\right] \quad \text{S18}$$

where $\Delta X_b$ and $\Delta X_w$ are the extension changes associated with the bound Ec$SSB_8$ and wrapped *Ec*$SSB_{17}$ states, respectively. Accordingly, we use $N_w$ to be 17 nt and the associated change in extension to be 0.08 nm/nt, as observed in Fig. 3B. The minimum extension change we observe at 50 nM (0.02 nm/nt) serves as an upper bound to the extension change associated with the bound *Ec*$SSB_8$ state. Therefore, we chose $N_b$ to be 8 nt (supported by AFM experiment, Fig. S4) and the associated extension change to be 0.015 nm/nt (supported by the monomeric *Ec*SSB mutant, Fig. S3). Using these experimental derived rates and amplitudes, the model correctly reproduces the time dependent binding curve (Fig. 2B) and equilibrium between the wrapped and unwrapped *Ec*SSB states (Fig. 2C) over the range of free protein concentrations observed. As the unwrapping rate of an *Ec*SSB on an unsaturated ssDNA ($k^0_{-w}$), is too slow to be measured in our system, we use $k^0_{-w}$= 0.01 $s^{-1}$ in the simulation that do not take stimulated unwrapping into account (Fig. 5C, dashed blue line), This estimation is based on the experimental observation that the maximally wrapped *Ec*SSB is stable over >100 s in the absence of free *Ec*SSBs in the solution at 12 pN Figs (1E, 3C).

#### *Supplemental Figures*

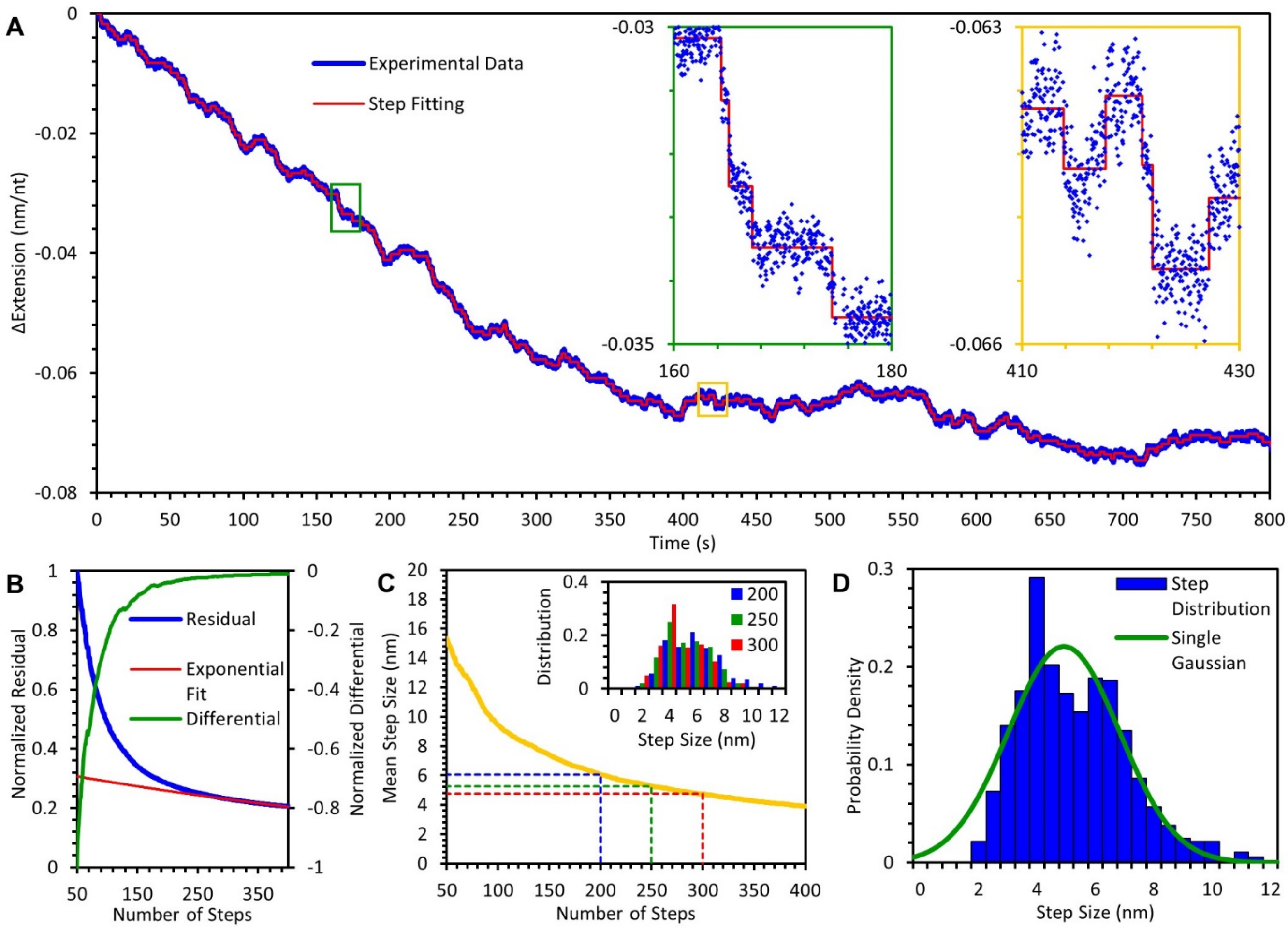


**Figure S1: Observations of single wrapping/unwrapping events at 12 pN.** (A) Extension-time profile of an ssDNA molecule incubated with 50 pM EcSSB (blue) is fitted to a step resolving function in MATLAB (red). The total number of steps is predefined (250 shown here), while the amplitude and timing of each step is determined by least square minimization. Initially decreasing steps are observed indicating *Ec*SSB wrapping events (green inset). As the system reaches equilibrium, steps in both directions are observed, indicating dynamic equilibrium of wrapping and unwrapping events (yellow inset). (B) The total residual of the step function fit (blue) decreases rapidly as the total number of fitted steps is increased. Past a certain threshold (~250 steps), the residual begins to decay at a slow exponential rate (red) with additional steps. The diminishing return in the goodness of fit can also be seen by plotting the differential of the residual (green). The fit rapidly improves in quality as the number of fitted steps approaches the number of real steps in the data (wrapping and unwrapping events). Beyond this point adding more steps simply fits noise in the system resulting in minor reduction of the residual. (C) The absolute value of the average step size (yellow) decreases as more steps are added to the fit but levels off when the correct number of steps are fit to the data. The inset shows that the distribution of individual step sizes peaks at ~5 nm for a wide range of fitted number of steps as determined by (B) (dashed blue, green, and red lines). (D) A representative distribution of individual step sizes is fitted to a Gaussian distribution that yields a mean step size of ~5 nM.

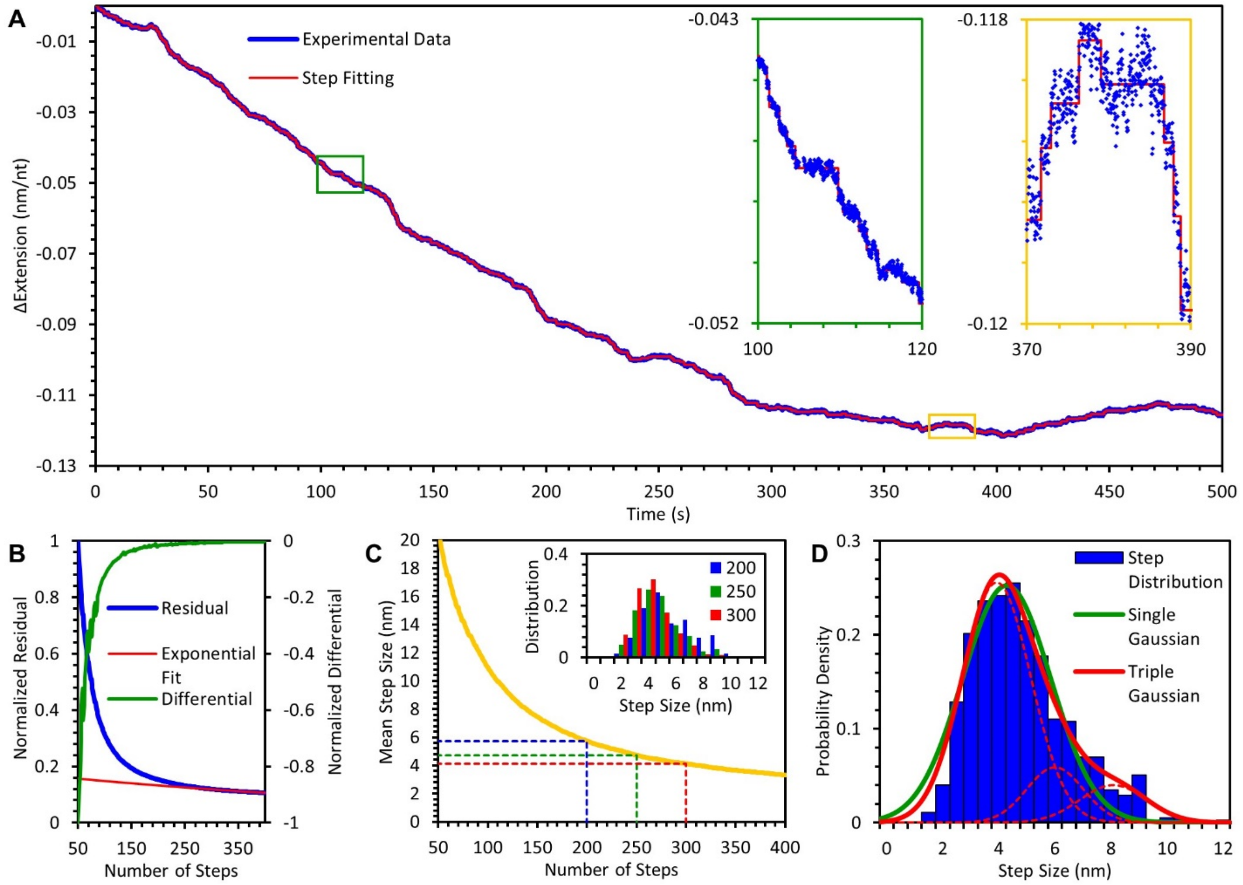


**Figure S2: Observations of single wrapping/unwrapping events at 7 pN.** (A) The analyses described in Fig. S1 here is repeated on an Extension-time profile obtained at 7 pN with 50 pM *Ec*SSB. Data is shown in blue and the step finding fit in red. Insets show steps that represent exclusively wrapping events, and wrapping events followed by unwrapping events, respectively. Panels (B)-(C) are the analysis described to legend in Fig. S1. (D) In contrast to that observed at 12 pN (Fig. S1), the distribution of the absolute step sizes (blue bars) is asymmetrical and cannot be fit by a single gaussian distribution. Instead, the step size distribution is fit with the sum of multiple gaussians, each representing a specific transition between the multiple wrapped states accessible to *Ec*SSB at 7 pN. The mean step sizes are estimated as ~4, ~6, and ~8 nm that likely represent transitions between *Ec*$SSB_{17}$, *Ec*$SSB_{35}$, and *Ec*$SSB_{56}$ modes.

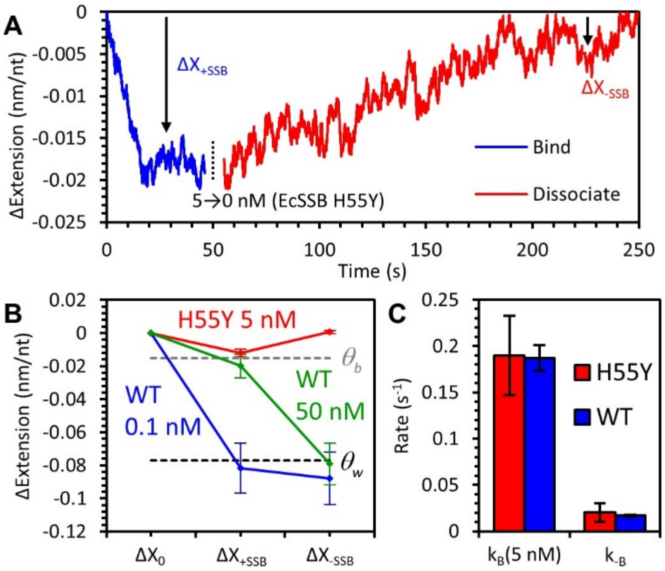


**Figure S3: Binding dynamics of monomeric *Ec*SSB (H55Y) to ssDNA.** (A) An ssDNA molecule held at 12 pN is incubated with 5 nM H55Y. At this concentration tetramer formation of H55Y is significantly compromised. In contrast to the binding of WT *Ec*SSB tetramer (Fig. 3), the initial change in ssDNA extension during H55Y binding (blue line) yields a much smaller amplitude with no secondary increase of extension as seen with the bind-unwrap process. Upon the removal of free H55Y (red line), the unbind-wrap process seen with the WT is also not observed. Instead the ssDNA extension slowly increases indicating direct dissociation events. Both the binding and dissociation curves are fit to single exponential functions to estimate the rates. (B) Average ssDNA extension changes after H55Y binding and dissociation. The ssDNA extension while bound by monomeric H55Y is consistent with that of the predicted bound but unwrapped $EcSSB_8$ state (Fig. 3). After dissociation, the ssDNA extension approaches its initial value, indicating dissociation of the H55Y. (C) Rates of H55Y binding and dissociation. The rate of binding ($k_B$) for 5 nM (monomer concentration) H55Y is the same as the rate of initial binding of an equivalent concentration of WT *Ec*SSB (1.25 nM tetramer concentration). The rate of H55Y dissociation is the same as the direct dissociation rate of WT *Ec*SSB at forces >20 pN that inhibit wrapping events.

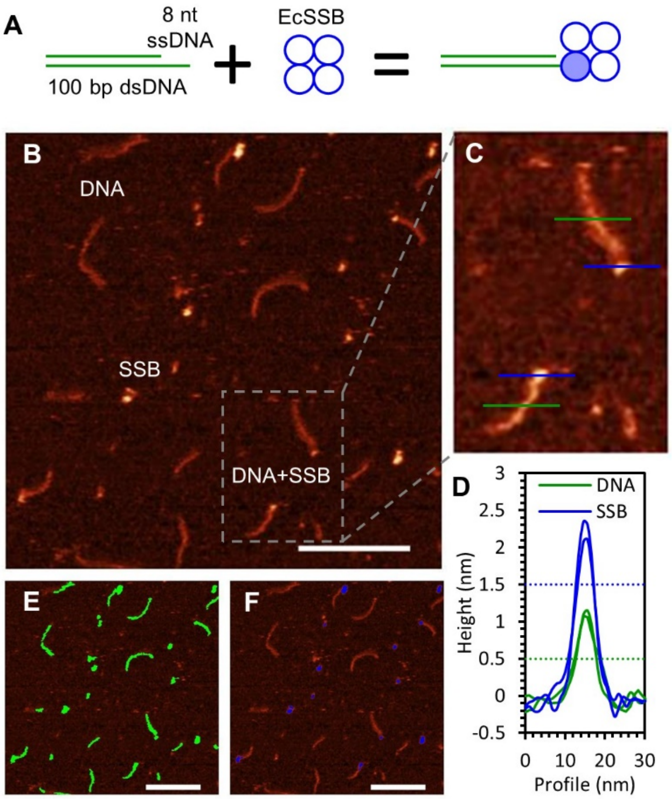


**Fig S4. AFM imaging of *Ec*SSB binding short ssDNA segments.** (A) A DNA construct consisting of 100 bp of dsDNA with an 8 nt poly(T) ssDNA overhang is incubated at equimolar concentrations (5 nM each). The schematic shows that the ssDNA overhang can only accommodate a single OB domain of *Ec*SSB. (B) These *Ec*SSB-DNA complexes are deposited on a treated mica surface and imaged using AFM (scale bar 100 nm). The DNA constructs appear as faint lines (ssDNA region cannot be observed distinctly from the dsDNA) and *Ec*SSB proteins appear as bright globules. *Ec*SSB tetramers localized at one end of the DNA construct indicate ssDNA binding. (C) Inset shows representative images of *Ec*SSB-DNA complexes. (D) Height profiles of dsDNA (green lines in (C)), and *Ec*SSB (blue lines in (C)), show that the measured maximum height of *Ec*SSB is approximately twice that of dsDNA. Therefore, a threshold of 0.5 nm (dotted green line), and 1.5 nm (dotted blue line) distinguish dsDNA, and *Ec*SSB, respectively, from the background. (E-F) Applying these thresholds to the entire image captures either both dsDNA and *Ec*SSB (green, threshold=0.5 nm) or *Ec*SSB only (blue, threshold=1.5 nm).

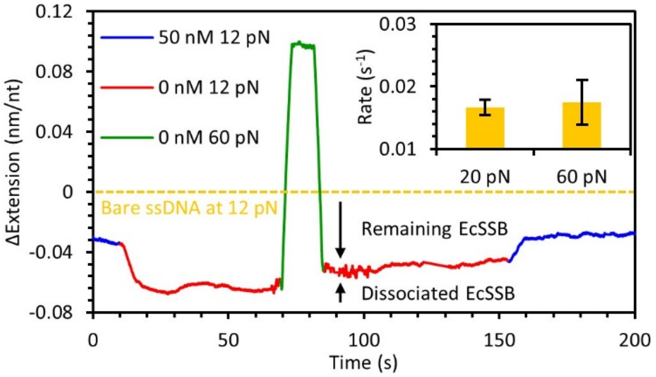


**Figure S5: Force-Jump experiments with *Ec*SSB and ssDNA**: After the *Ec*SSB-ssDNA complex is saturated upon initial incubation (blue, bind-wrap transition is not shown), and the free protein is removed (unbind-wrap shown in red) at 12 pN, the *Ec*SSB-ssDNA complex equilibrates at the maximally wrapped state with no dissociation observed. Then the applied force on the template is rapidly increased, held at 60 pN and returned to 12 pN during a ~15 s timescale. Most protein remains bound during this transition and is able to rewrap the ssDNA at 12 pN, as evidenced by the mostly preserved change in extension (with respect to protein-free ssDNA). Additionally, reintroducing free protein (at t~150 s), results in an increase in extension that is consistent with the rebind-unwrap process (Fig. 3), rather than a biphasic binding curve observed with *Ec*SSB binding to protein-free ssDNA, reinforcing that most *Ec*SSB remains bound. The change in ssDNA extension before and after the 60 pN force spike is used to estimate the amount of protein that dissociated at high force and calculate the rate of *Ec*SSB dissociation in the absence of wrapping events ($k_{-b}$). This value (inset) is equal to the rate of dissociation we directly observe at 20 pN.

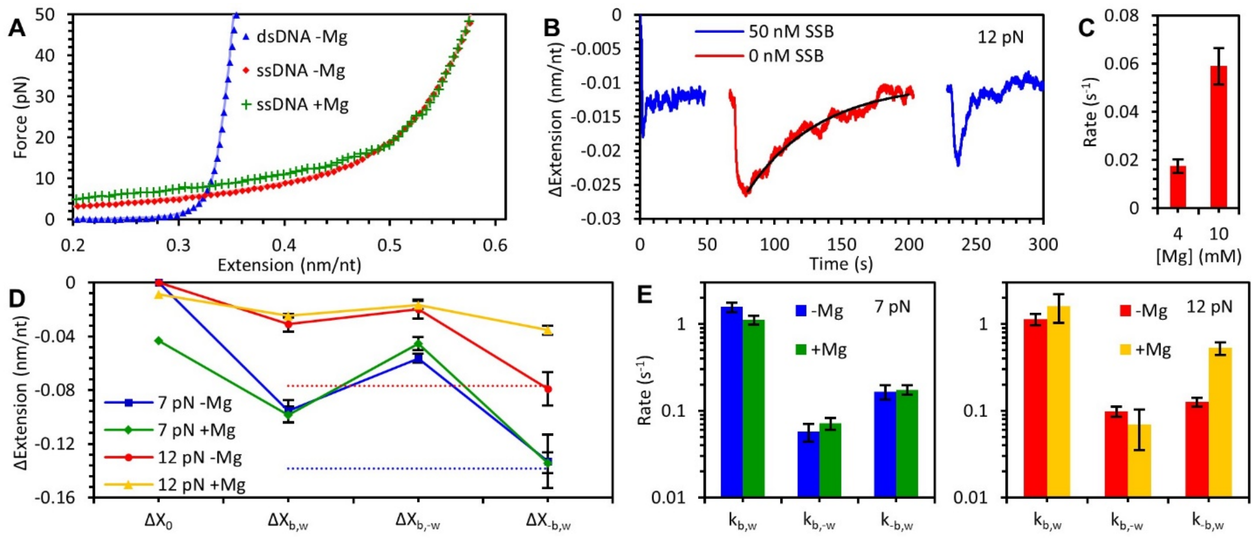


**Figure S6: Impact of Magnesium on *Ec*SSB binding dynamics**. (A) Magnesium (4mM $Mg^{2+}$) stabilizes the local secondary structures on ssDNA (such as hairpins), resulting in apparent reduction in extended length (green) at forces <20 pN. At forces > 20 pN the secondary structures are eliminated rendering similar ssDNA extension profiles in the presence (red) and absence of $Mg^{2+}$. (B) In the presence of $Mg^{2+}$, *Ec*SSB still exhibits a biphasic binding pattern (blue). After removing the free *Ec*SSB from the solution, the unbind-wrap process is followed by direct dissociation at 12 pN. This is similar to what is observed at 20 pN in the absence of $Mg^{2+}$ (Fig. 4). (C) The rate of the direct dissociation measured here increases as the $Mg^{2+}$ concentration is increased. (D) The total change in extension due to *Ec*SSB binding is similar both in the presence and absence of magnesium. Note that the length decrease of the protein-free ssDNA due secondary structures in the presence of $Mg^{2+}$ is taken into account. (D) The rates associated with each phase of the *Ec*SSB binding (as described in Fig. 3) are similar in the presence and absence of $Mg^{2+}$. Overall the results suggest that the presence of $Mg^{2+}$ destabilizes the low order *Ec*SSB wrapped conformations, and enhances *Ec*SSB dissociation from the unsaturated complex when wrapping is unfavorable due to template tension.
